# On the Ca^2+^ elevation in vascular endothelial cells due to inositol trisphosphate-sensitive store receptor activation: a data-driven modeling approach

**DOI:** 10.1101/2022.10.17.512592

**Authors:** Alberto Coccarelli, Sanjay Pant

## Abstract

Agonist-induced Ca^2+^ signalling is essential for the regulation of many vital functions in endothelial cells (ECs). A broad range of stimuli elevate the cytosolic Ca^2+^ concentration by promoting a pathway mediated by inositol 1,4,5 trisphosphate (IP_3_) which causes Ca^2+^ release from intracellular stores. Despite its importance, there are very few studies focusing on the quantification of such dynamics. Here, by using data from isolated ECs, we constructed and tested a minimalistic model that can be used for capturing the main features (averaged over a cell population) of the Ca^2+^ response to different IP_3_ stimulation levels. A suitable description of Ca^2+^-regulatory function of inositol 1,4,5 trisphosphate receptors (IP_3_Rs) is identified by comparing the different model variants against experimental mean population data. The same approach is used to numerically assess the relevance of cytosolic Ca^2+^ buffering, as well as Ca^2+^ store IP_3_-sensitivity in the overall cell dynamics. The variability in the dynamics’ features observed across the population can be explained (at least in part) through variation of certain model parameters (such as buffering capacity or Ca^2+^ store sensitivity to IP_3_). The results, in terms of experimental fitting and validation, support the proposed minimalistic model as a reference framework for the quantification of the EC Ca^2+^ dynamics induced by IP_3_Rs activation.

## 1 Introduction

Among the myriad of pathways regulating vascular function, Ca^2+^ release from intracellular stores plays a fundamental role in the control of blood pressure and flow across the whole circulation [1, 2]. The associated Ca^2+^ elevation in the endothelial cytosol has indeed manifold effects on tone regulation including the enhancement of nitric oxide synthase (eNOS) activity [3] and potassium (SKCa/IKCa) channels activation which leads to the hyperpolarization of the plasma membrane [4].

Despite its involvement in different agonist stimulations [5, 6], the contribution of Ryanodine Receptors (RyRs) located in the sarcoplasmic/endoplasmic reticulum (SR/ER) membrane to EC function is still not clear as their activation through caffeine does not seem to elevate cytosolic Ca^2+^[7, 8]. On the other hand, Ca^2+^ release from endoplasmic reticulum (ER) due to activation of IP_3_Rs plays a prominent role in several exogenous ligand-induced pathways. Among these, the Ca^2+^ signalling induced by muscarinic cholinergic receptor M3 stimulation has been ascertained across various experimental settings [9, 10, 11, 12]. Activation of muscarinic M3 receptors with acetylcholine (ACh) enhances the production rate of diacylglycerol and IP_3_, which is then followed by a cytosolic Ca^2+^ elevation. The initial increase in cytosolic Ca^2+^ is caused by IP_3_ binding to its receptor on ER which enables a store IP_3_-mediated Ca^2+^ release. The activation of a single IP_3_R on ER initiates the store Ca^2+^ release as a localized Ca^2+^ blip, which may expand to neighbouring IP_3_Rs generating a wider signal, defined as Ca^2+^ puff. For rising IP_3_ concentration levels, this signal can augment in frequency and propagate to other IP_3_R clusters. The increase in Ca^2+^ nearby the IP_3_R favours the ER Ca^2+^ release through a positive feedback mechanism, defined as Ca^2+^-induced Ca^2+^-release (CICR) mechanism. Depending on the intracellular conditions and stimulation level, this signal may further propagate causing a transient global cell elevation in Ca^2+^, denoted as Ca^2+^ wave, that can activate different types of membrane channels [13]. Whilst RyRs in the SR generate local Ca^2+^ signalling events (denoted as ‘sparks’) allowing the communication with the juxtaposed plasma membrane, IP_3_Rs are capable of generating spatially fixed local release events (called ‘pulsars’) which project through the elastic lamina to the neighbouring smooth muscles membranes [14]. In pressurized mesenteric arteries, spontaneous Ca^2+^ events appear to originate from ER IP_3_Rs even under resting conditions, and these appear to be regulated by surrounding smooth muscle cells via myoendothelial gap junctions [15]. The depletion of Ca^2+^ in the intracellular stores triggers Ca^2+^ influx through an capacitative entry mechanism (CCE) that serves for sustaining the cytosolic Ca^2+^ elevation and ultimately enables ER Ca^2+^ refilling [16, 17, 3]. This store-operated effect involves the activation of transient receptor potential (TRP) channels located on the EC plasma membrane, whose activation under agonist intervention was characterized in different studies [18, 19, 20, 21, 22, 23, 24]. Once the agonist effect vanishes, Ca^2+^ extrusion and re-sequestration through Na^+^-Ca^2+^ exchanger (NCX) and Ca^2+^ ATPase pumps (PMCA) retake the cytosolic Ca^2+^ concentration to its original basal level [25, 26].

To operate as an effective sensor, the endothelium is endowed with different cell phenotypes [27], which may exhibit remarkably heterogeneous Ca^2+^ responses to agents involving the IP_3_R pathway such as ACh [28, 13]. However, Carter and Ogden [29] showed that the dependency between the maximum IP_3_Rs Ca^2+^ flux and cytosolic IP_3_ concentration across a porcine aortic EC population can be well captured through an activation function which is regulated by the Ca^2+^ concentration. Therefore the EC discrepancy in the response to ACh may not only be explained with different expression levels of in IP_3_R isoforms, but also due to other cellular components involved in Ca^2+^ buffering, extrusion and store sequestration. Furthermore, this study well documented the inhibitory role of Ca^2+^ on IP_3_Rs by considering a broad range (up to saturation level) of IP_3_ stimulations and showed that IP_3_Rs inhibition occurs independently of the level of IP_3_ stimulation. It is also worth noting that, apart from the study by Carter and Ogden [29], there is scarce quantitative information in the literature about the activation role of Ca^2+^ on EC IP_3_Rs at low cytosolic Ca^2+^ concentrations.

Depending on the level of detail required, Ca^2+^ release via ER IP_3_R can be modelled by using different approaches including Markov models, gating models and modal models [30]. De Young-Keizer model developed a framework [31, 32, 33] for describing the kinetics of the IP_3_R channel release in the ER, which provided the basis for the development of stochastic models for studying CICR and Ca^2+^ propagation phenomena (from puffs to waves) within the cytosol [34, 35, 36, 37, 38]. On the other hand, gating models provide a minimalistic description of the Ca^2+^ release due to IP_3_Rs activation which is particularly suitable for the incorporation into whole-cell dynamics models [39, 40, 30].

With respect to vascular ECs, some relevant modelling studies were carried out towards the characterization of endothelial Ca^2+^ response to agonist stimulation. Wiesner et al. [41] proposed a mathematical model for describing the calcium dynamics originating in endothelial after thrombin intervention. This work provided a detailed description of the kinetics involved from the membrane receptor to the induced Ca^2+^ release. The modelling of the IP_3_R channels followed a previous study carried out on rat basophilic leukemia cells [42]. The work by Wiesner al. laid the foundation for the development of other models [43, 44, 45, 46], which were used for investigating the effects of different agonists and shear stress on the EC Ca^2+^ dynamics. Since the activation of IP_3_Rs by exogenous agonists can only occur via signalling mediators in the cytosol, the predictive capacity of the corresponding model is hindered by uncertainties associated with the multiple kinetics involved.

To the best of our knowledge, there is no established modelling framework that can quantitatively describe the endothelial Ca^2+^ dynamics following the IP_3_-induced Ca^2+^ release from ER. In this work, we leverage on the Ca^2+^ dynamics data obtained by direct stimulation of IP_3_Rs by Carter and Ogden [29], which allow us to exclude the uncertainties associated with the kinetics regulating the ligand-induced IP_3_ cytosolic production from the model. Despite the heterogeneous behaviour observed in different stimulations involving the IP_3_R pathways [29, 28, 13], we hypothesize the existence of a ‘reference’ model able to capture the mean Ca^2+^ transient features across an EC population. We test this hypothesis by assessing the performance (in terms of fitting and validation) of a novel, minimalistic Ca^2+^ dynamics model (alongside some more complex variants) which uses a gating function for regulating the IP_3_-induced Ca^2+^ release in the cytosol. We then further speculate that cellular variability may be explained as deviation in some of its parameters defining cellular functions such as Ca^2+^ sequestration and cytosolic buffering. The identified model will ultimately enable us to the estimate physical quantities which are difficult to measure directly from the experiment.

## 2 Methods

### 2.1 Mathematical model

#### 2.1.1 Experimental conditions and stimulation

The mathematical model is defined by considering the experimental conditions adopted by Carter and Ogden [29]. In brief, isolated ECs from porcine thoracic aorta were stimulated with different concentration of IP_3_ in whole cell patch-clamp settings (holding membrane potential of 0 mV). Since the EC holding membrane potential was 0 mV, the contribution of transmembrane ionic currents driven by the potential gradient can be ignored in the cytosolic Ca^2+^ concentration balance (membrane hyperpolarization elevates the cytosolic Ca^2+^ by increasing Ca^2+^ influx, as shown in [29]). Caged IP_3_ (P-4 and P-5 isomers) was released in the cytosol by means of photolysis lasting ~ 1 ms. Therefore the cell stimulation consisted in a pulse increase of the average cytosolic IP_3_ concentration (*i*), whose dynamics can be modelled with a first-order kinetics reaction equation:

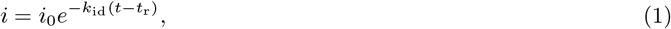

where *i*_0_ is the initial cytosolic IP_3_ concentration after 1 ms pulse of near UV light, *k*_id_ is the IP_3_ dissociation constant due to the interaction with other cytosolic components, and *t*_r_ is the release time.

#### 2.1.2 Ca^2+^ store release

For the considered dynamics, the variation in free Ca^2+^ concentration across the cytosol is not considered significant and therefore the currents between the cytosol and the ER and the extracellular space can be expressed as function of the average free cytosolic Ca^2+^ concentration (*c*_c_). For the sake of brevity, from here onwards we omit the ‘free’ in front of cytosolic Ca^2+^, whilst the bounded cytosolic Ca^2+^ is indicated as ‘buffered’ cytosolic Ca^2+^. The Ca^2+^ dynamics triggered by the activation of the IP_3_-dependent store channels is the result of the simultaneous interaction between different channels, pumps and exchanger expressed at membrane, cytosolic and ER levels. Here we capture the key characteristics of IP_3_R-mediated Ca^2+^ release by employing a minimalistic approach which incorporates a gating variable. The endogenous Ca^2+^ release flux from ER due to IP_3_Rs activation (*J*_ef_, see Figure 1) is made dependent on the cytosolic levels of IP_3_ and Ca^2+^ through the following expressions [39, 40, 30]

**Figure 1:**
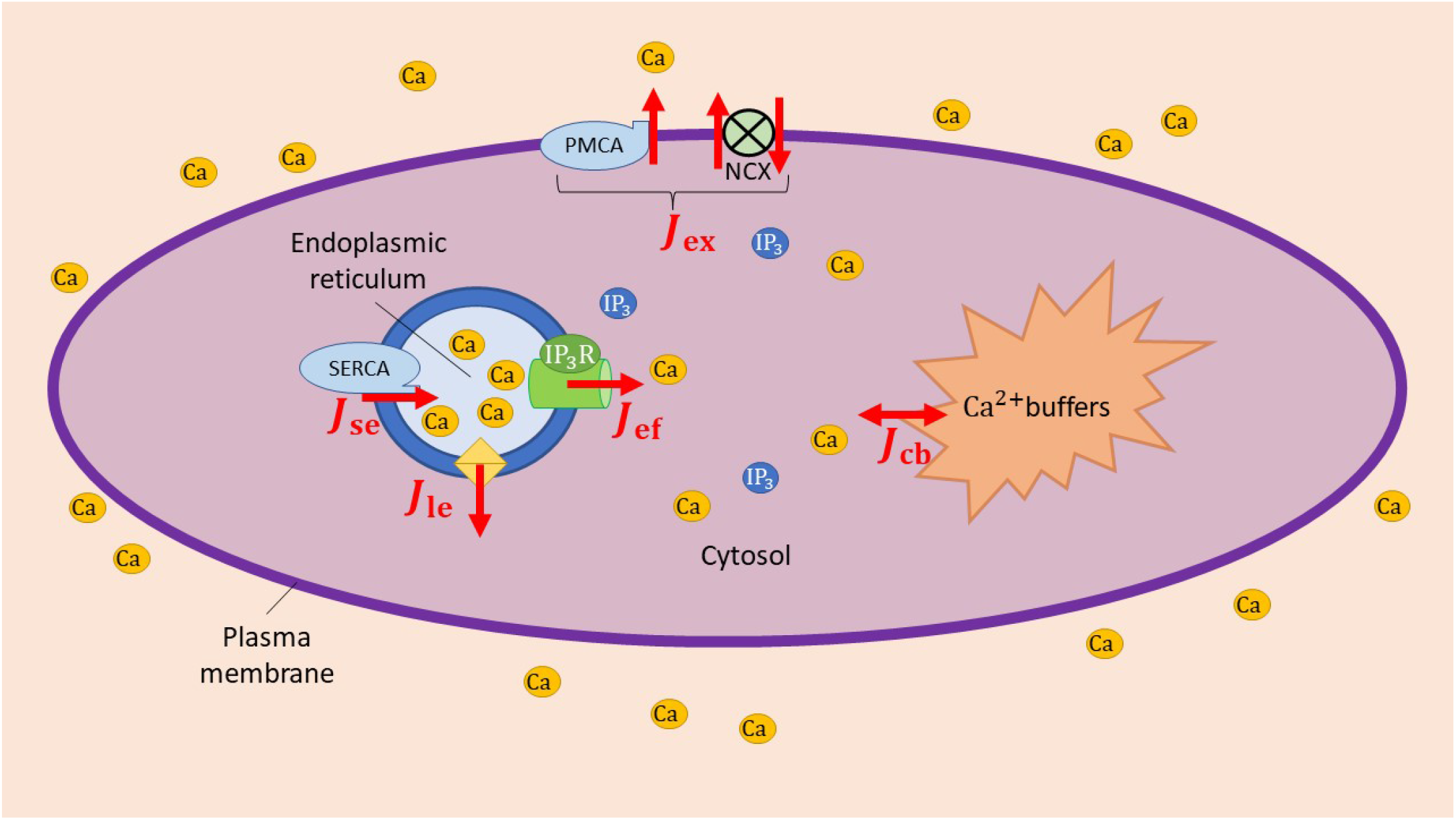
Key components of the cellular Ca^2+^ dynamics model. *J*_ef_: ER IP_3_-sensitive Ca^2+^ release flux; *J*_le_: ER Ca^2+^ leakage flux to cytosol; *J*_se_: ER Ca^2+^ sequestration flux; *J*_ex_: Ca^2+^ extrusion flux from cytosol; *J*_bc_: Ca^2+^ buffering flux.

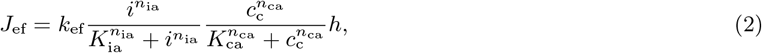

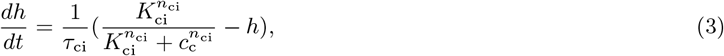

where *h* is a gating variable representing the availability/state of IP_3_ receptors, *k*_ef_ is the nominal ionic flux, *τ*_ci_ is the time constant governing the channel inactivation, whist *K*_ia_, *K*_ca_, *K*_ci_ and exponents *n*_ia_, *n*_ca_ and *n*_ci_ are the coefficients of the corresponding Ca^2+^-dependent activation and inactivation functions. Here, due to the lack of experimental evidence, we assume that *K*_ci_ and *τ*_ci_ are *i*-independent and *c*_c_-independent, respectively. Eq. (2) implies assuming the interaction between IP_3_ and IP_3_Rs instantaneous [47], and that the Ca^2+^ release flux is proportional to the fraction of open channels, which is inline with previous work [41]. In the considered experiment, the time-length associated with the Ca^2+^ store release is extremely short (~ 1 s) and therefore it is reasonable to assume that the variation in Ca^2+^ concentration in the ER (*c_s_*) is minimal and does not impact the Ca^2+^ current from ER. Consequently, the membrane Ca^2+^ influx associated with store-depletion can also be considered negligible. We highlight that the IP_3_-mediated Ca^2+^ release is not made proportional to the difference in average Ca^2+^ concentration between the store and the cytosol because the latter is not necessarily representative of the Ca^2+^ gradient nearby the channels. The Ca^2+^ leakage from the ER to the cytosol (see Figure 1) is accounted for as

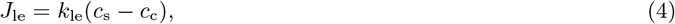

where *k*_le_ indicates the permeability of the ER membrane while *c*_s_, as explained above, is assumed to be constant.

#### 2.1.3 Ca^2+^ sequestration, extrusion and buffering

Ca^2+^ re-uptake into ER (*J*_se_) occurs via SERCA pump while Ca^2+^ extrusion from the cytosol (*J*_ex_) is operated by membrane proteins such as PMCA and NCX (see Figure 1). These can generally be described as Ca^2+^-activation functions [41, 46, 30]

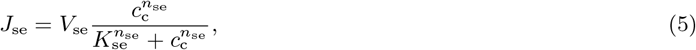

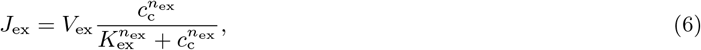

where *V*_se_ and *V*_ex_ are the nominal fluxes and *K*_se_, *K*_ex_, *n*_se_ and *n*_ex_ are the activation function coefficients corresponding to the Ca^2+^ sequestration and extrusion from the cytosol. More complex formulations for these pumps and exchangers can be found in literature [46, 30]. Despite such pumps and exchangers playing a physiologically essential role in the Ca^2+^ depletion from the cytosol, in the considered experiment their contribution to the cytosolic Ca^2+^ balance is limited [29]. In ECs, the Ca^2+^ extraction rate of the membrane pumps remains low also because their characteristic time for full activation is considerably higher than the considered time-frame [25, 48]. Due to this, together with the fact that the functioning of such proteins in ECs is poorly characterized and distinguished in literature, we assume that all Ca^2+^ removal mechanisms from the cytosol can be incorporated together under only one flux, whose activation depends solely on cytosolic Ca^2+^ concentration (*c*_c_). To avoid introducing new notation, we refer to this contribution with the flux *J*_se_ introduced in Eq. (5) whilst the flux *J*_ex_ introduced in Eq. (6) will be considered negligible for the cytosolic Ca^2+^ balance.

Different types of intracellular buffers, such as calmodulin and fluorescence dye, act by dampening the variation in free cytosolic Ca^2+^ concentration. There are different ways for including this effect into the model. A classical approach treats the (averaged in space) buffered cytosolic Ca^2+^ concentration (*c*_b_) as an independent variable, whose rate of change is determined via [41, 46]

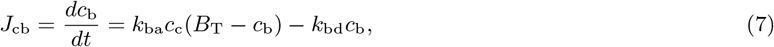

where *k*_ba_ and *k*_bd_ are, respectively, the association and dissociation rate constants for Ca^2+^ binding to cytosolic proteins whilst *B*_T_ is the concentration of the total buffering proteins present in the cytosol. However, in the considered experiments [29], the variation in cytosolic free Ca^2+^ appeared proportional to the change in the total (free plus buffered) cytosolic Ca^2+^. Therefore, Ca^2+^ buffering may be also approximated as an instantaneous process and simply accounted for by means of a parameter *β*_cb_ representing the fraction of non-buffered Ca^2+^ with respect to the total. In this case, the rate of change in the cytosolic Ca^2+^ concentration can be expressed as

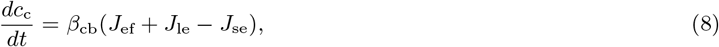

where *β*_cb_ can be assumed to be constant or, alternatively, defined through the following expression [49, 50]

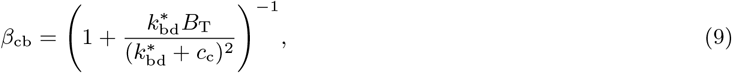

in which 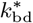 is the corresponding Ca^2+^ dissociation rate constant. If the Ca^2+^ buffering is instead treated as a dynamic process, the ionic mass balance yields

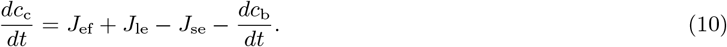

#### 2.1.4 Solving the Ca^2+^ dynamics

The solution of the Ca^2+^ dynamics can be obtained, depending on the adopted Ca^2+^ buffering approach, by solving either the system of Eq. (3) and (8) for the variables *h* and *c_c_* or the system of Eq. (3), (7) and (10) for the variables *h*, *c*_c_ and *c*_b_. The time evolution of IP_3_ concentration *i* is considered independent from cytosolic Ca^2+^ concentration and therefore Eq. (1) is computed before the Ca^2+^ dynamics. In this study we consider five Ca^2+^ dynamics model variants, each of them governed by a set of parameters to be determined (see Table 1). The system of ODEs representing the Ca^2+^ dynamics is integrated over time by using the function ‘odeint’ from the python library NumPy, with time-step Δt=0.001 s (this value was set after appropriate time-convergence analysis).

**Table 1:**
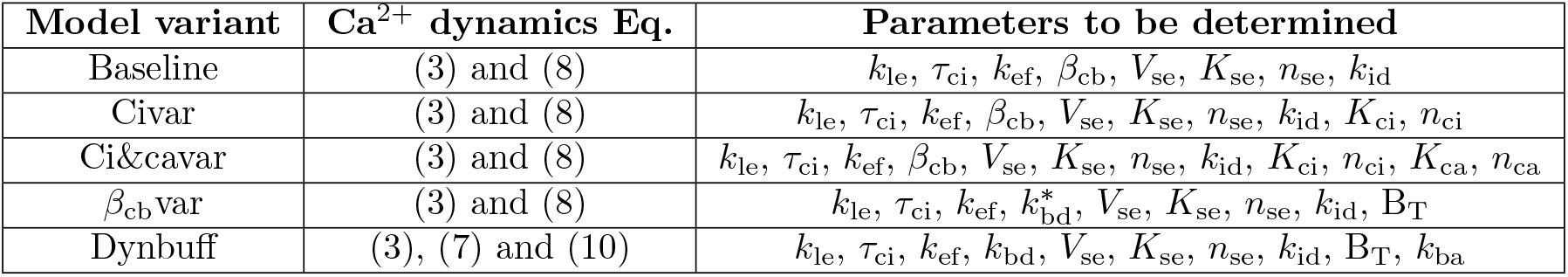
Model variants for representing Ca^2+^ dynamics in EC. For each model variant, the fixed parameters are reported in Section 3.

**Table 2:**
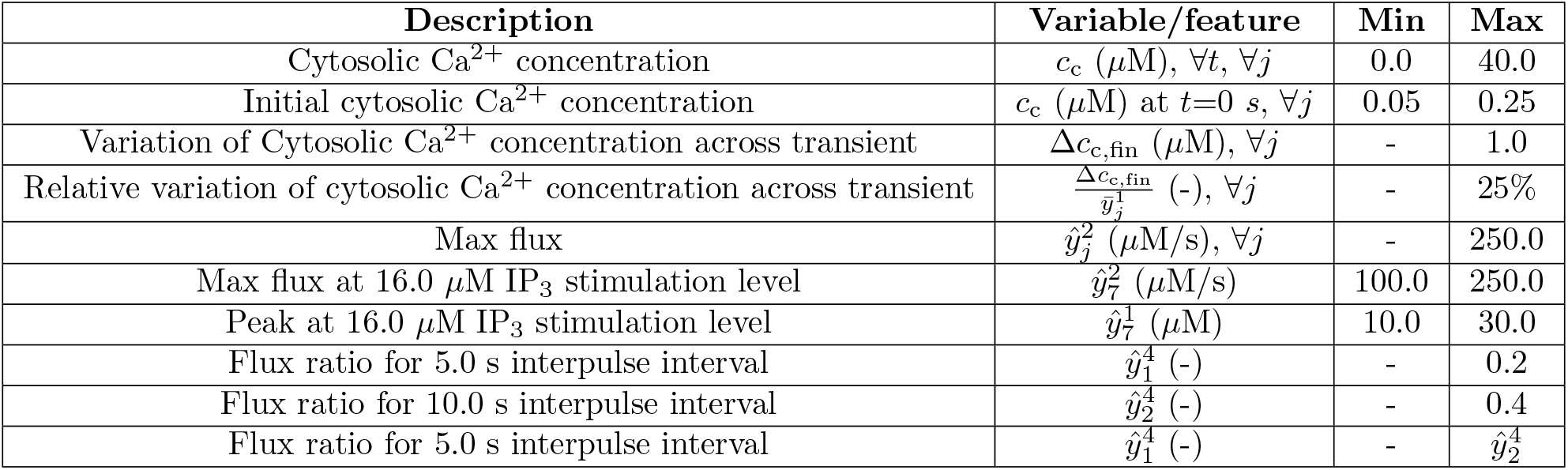
Imposed constraints in the optimization procedure. Δ*c*_c,fin_ is the difference in cytosolic Ca^2+^ concentration between the final (*t* > 20.0 s) and initial time step. ∀*t*: at any time; ∀ *j*: for any IP_3_ stimulation level.

### 2.2 Model parameter identification

Here we present the methodology for identifying the set of parameters for each model variant that best fit the mean EC population features. We devise an optimization problem in which the solution is the set of model parameters that minimizes the discrepancy (quantified by a cost function) between the features of the simulated Ca^2+^ transient and the corresponding measurements across an EC population. A set of model parameters is accepted only if the error (see Section 2.2.1) with respect to the experimental mean population data is below a certain ‘ad-hoc’ threshold. Four features of the Ca^2+^ concentration transient are considered for the fitting: the ‘Peak’, ‘Max flux’, ‘Termination rate per unit of flux’ (see Figure 2) and ‘Flux ratio’ obtained from two consecutive stimulations with interpulse interval Δ*t_inter_* (see Figure 4 in [29]). The first three features were recorded across the IP_3_ stimulation range 0.2–16.0 *μ*M, whilst the latter feature was evaluated with a first *i* impulse at 0.8 *μ*M, followed by a second impulse at 0.7 *μ*M, as documented in [29]. The feature Flux ratio can be numerically evaluated as ratio between the values of the gating variable *h* obtained immediately (1 time-step Δt) after the two consecutive stimulations (separated by a interpulse Δ*t_inter_*). According to the experimental study, 16.0 *μ*M of IP_3_ is high enough to saturate all the corresponding store receptors, and therefore no IP_3_ stimulation beyond such level is considered. Details on the feature extraction from Ca^2+^ concentration time-recordings are reported in the Appendix.

**Figure 2:**
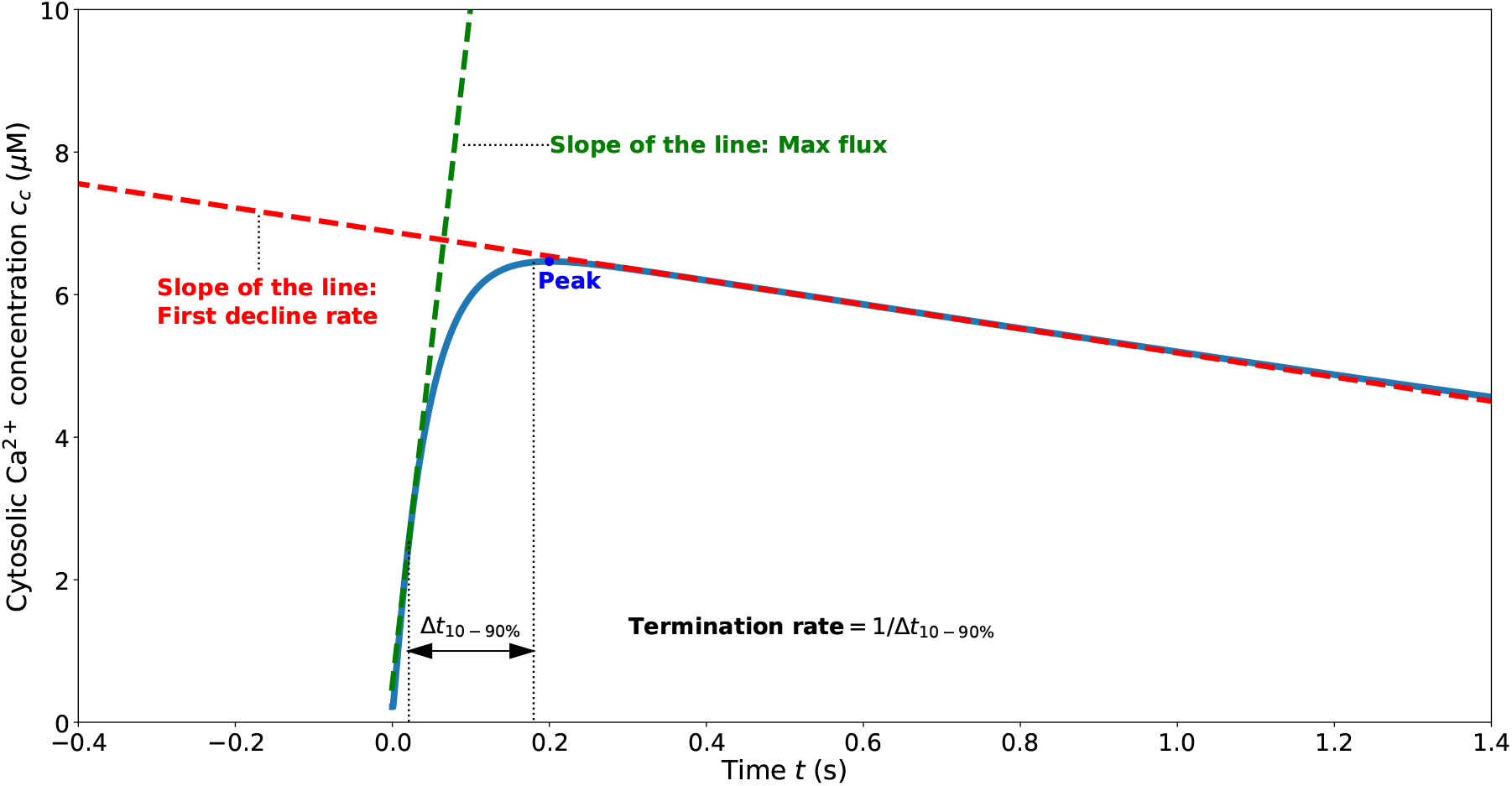
Features measured from EC cytosolic Ca^2+^ concentration transient. The Termination rate per unit of flux is evaluated by dividing the Max flux by the ‘Termination rate’, with the latter being the inverse of 10–90 % rise time (Δ*t*_10−90%_).

#### 2.2.1 Cost function

The cost function 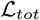 to be minimized is made of two components 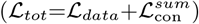. 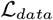 is the sum of all relative errors of the simulated features (ŷ) with respect to average values from experimental data (ȳ):

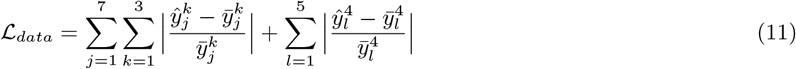

where *j* indicates an IP_3_ stimulation level *i*_0_ = {0.2, 0.4, 0.8, 1.4, 2.0, 5.0, 16.0 *μ*M}, *k* corresponds to a Ca^2+^ transient feature (Peak, Max flux, Termination rate per unit of flux, Flux ratio) and *l* identifies a interpulse interval Δ*t_inter_* = {5.0, 10.0, 15.0, 20.0, 25.0 s}.

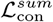 accounts for the physical constraints that the Ca^2+^ transient trace must satisfy. These are introduced to refine and accelerate the search by discarding solutions which do not lie within the expected physical value ranges. Each constraint contribution is included into the cost function as an individual penalty 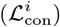:

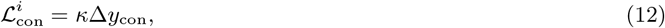

where *κ* is the penalty parameter (1.0e6) and Δ*y*_con_ defines the distance of the variable/feature with respect to the allowed interval boundary. Set of parameters that yield a ‘First decline rate’ (see Figure 2) lower than −5.0 *μ*M/s are also discarded in the optimization process. According to measurements (see Figure 3 in [29]), this latter feature is sparse with respect to Peak, and therefore is used only for discarding undesirable solutions.

**Figure 3:**
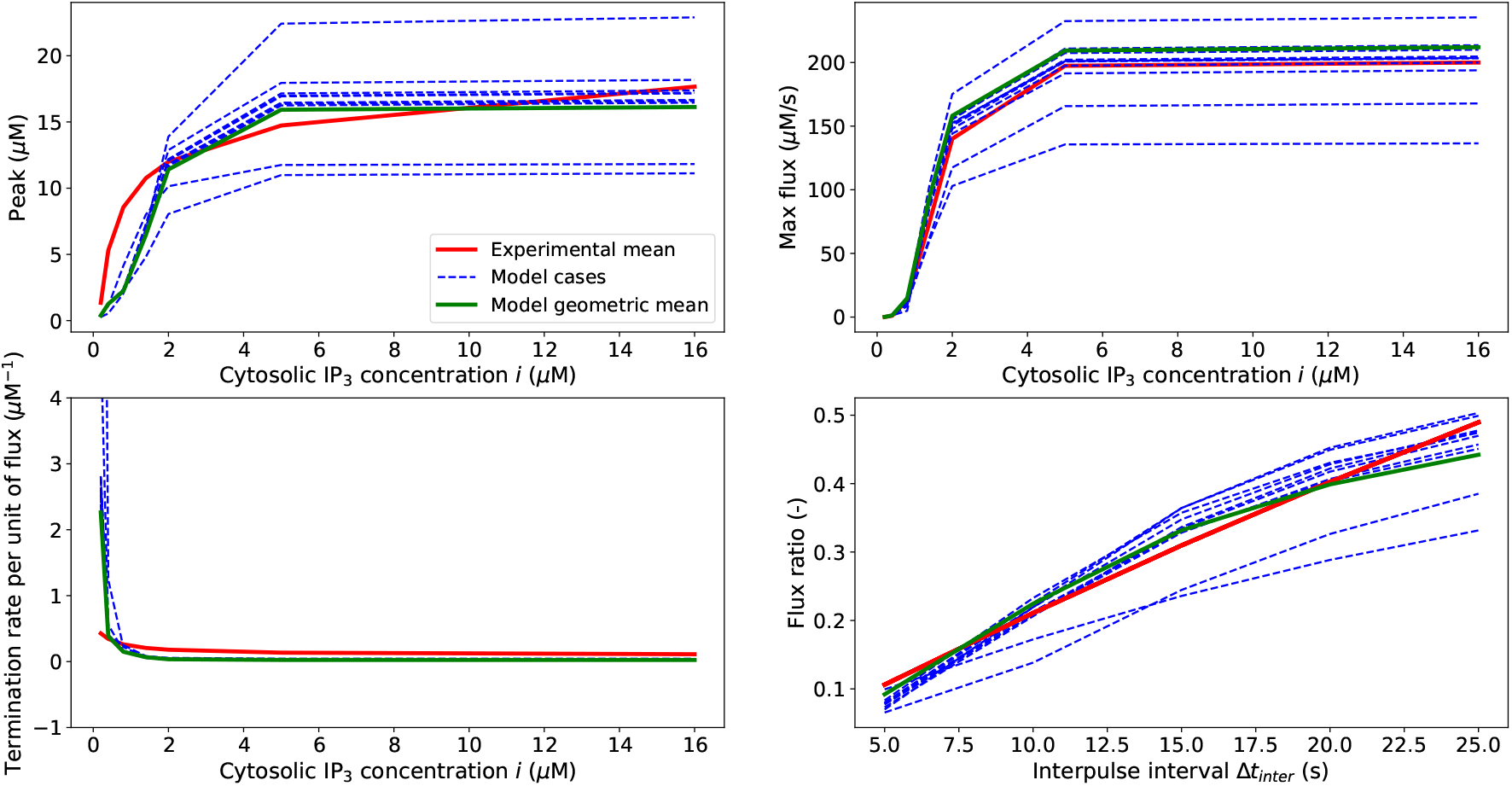
Comparison between the experimental mean against 10 simulated cases and geometric mean of the Baseline model.

#### 2.2.2 Optimization algorithm

The parameter set is identified by minimizing the cost function through the covariance matrix adaptation evolution strategy (CMA-ES) [51, 52]. The search stops when either the cost function goes below a ‘ad-hoc’ threshold (chosen to be 3.5 after empirical evaluation) or the number of iterations employed by the search-algorithm go above a pre-set maximum number of iterations (500). Given the high number of model parameters and due to the stochastic nature of the search algorithm, the optimizer may lead to different optimal set of parameters for the same model variant. Therefore, for any model variant, multiple cases (1,000) are used for identifying the most likely space of parameters. Since any considered model parameter (*ξ*) can be only positive, we express it as *ξ* = *ξ*_0_*e^α^*, where *ξ*_0_ is a constant (kept fixed during the optimization process) whilst *α* is the term through which the parameter *ξ* is varied. At the start of the search *α* is set to zero for each parameter and then updated by the optimization algorithm.

## 3 Results

Here we target the identification of the parametric space for each model variant, and the assessment of their performance by employing the quantitative experimental data from the study Carter and Ogden [29]. For doing this, different model variants are considered (introduced from the simplest to the most complex). This allows to identify the model components which are essential for capturing the experimentally-observed EC behaviour. The model is then used for making predictions on how EC variability may impact the IP_3_-induced Ca^2+^ dynamics.

### 3.1 Baseline model

We start by considering the simplest model, defined as ‘Baseline’ model for which Ca^2+^ buffering can be approximated as an instantaneous and Ca^2+^-independent process (*β*_cb_ is constant). Furthermore, for this model, we assume that Ca^2+^ affects the IP_3_Rs in the same manner as reported in [46] (we fix *K*_ci_=1.0 *μ*M, *n*_ci_=3.8, *K*_ca_=0.0 *μ*M). The coefficients for the *i*-dependent activation function of the ER Ca^2+^ flux are directly taken from [29] (we fix *K*_ia_=1.6 *μ*M, *n*_ia_=3.8). As anticipated earlier, the Ca^2+^ concentration in the ER (*c*_s_) can be considered constant, equal to 3,000 *μ*M. The model parameters (*k*_le_, *τ*_ci_, *k*_ef_, *β*_cb_, *V*_se_, *K*_se_, *n*_se_, *k*_id_) are identified by solving the optimization problem described in Section 2.2. Around 15 % of the cases were discarded because their cost function did not satisfy the imposed error threshold within the maximum number of iterations of the optimizer (500).

Figure 3 compares the curves of 10 different model parameter sets (obtained from 10 solutions of the optimization problem) and the solution obtained from the model with the geometric mean of (all) the parameters against the mean population experimental values. The geometric mean (for a given model variant) is obtained by averaging the variable term of the parameter *α* (and not the final parameter value *ξ*) across all the parameter sets obtained as solution of the optimization problem. Table 3 reports the initial/guess value and geometric mean for each parameter of the Baseline model, obtained by considering 847 cases/solutions.

**Table 3:** Initial parameter set and geometric mean of the parameter set cases for the Baseline model.

| - | $k_{le}$ (s<sup>-1</sup>) | $\tau_{ci}$ (s) | $k_{ef}$ (μM/s) | $\beta_{cb}$ (-) | $V_{se}$ (μM/s) | $K_{se}$ (μM) | $n_{se}$ (-) | $k_{id}$ (s<sup>-1</sup>) |
| --- | --- | --- | --- | --- | --- | --- | --- | --- |
| Initial ($\xi_0$) | 1.0e-6 | 2.0 | 2.0e2 | 1.3e-1 | 5.0 | 1.5e-1 | 2.0 | 1.0 |
| Geometric mean | 4.4e-7 | 7.378e-2 | 1.585e3 | 1.882e-1 | 1.935e1 | 3.628 | 3.579 | 2.207e-1 |

As anticipated, the adopted optimization procedure leads to the identification of the parameter space for the Baseline model by comparing the simulated EC dynamics due to IP_3_Rs activation against the mean response features extracted from the cell population. This optimization problem yields multiple solutions as reflected in the parameter space reported in Figure 4.

**Figure 4:**
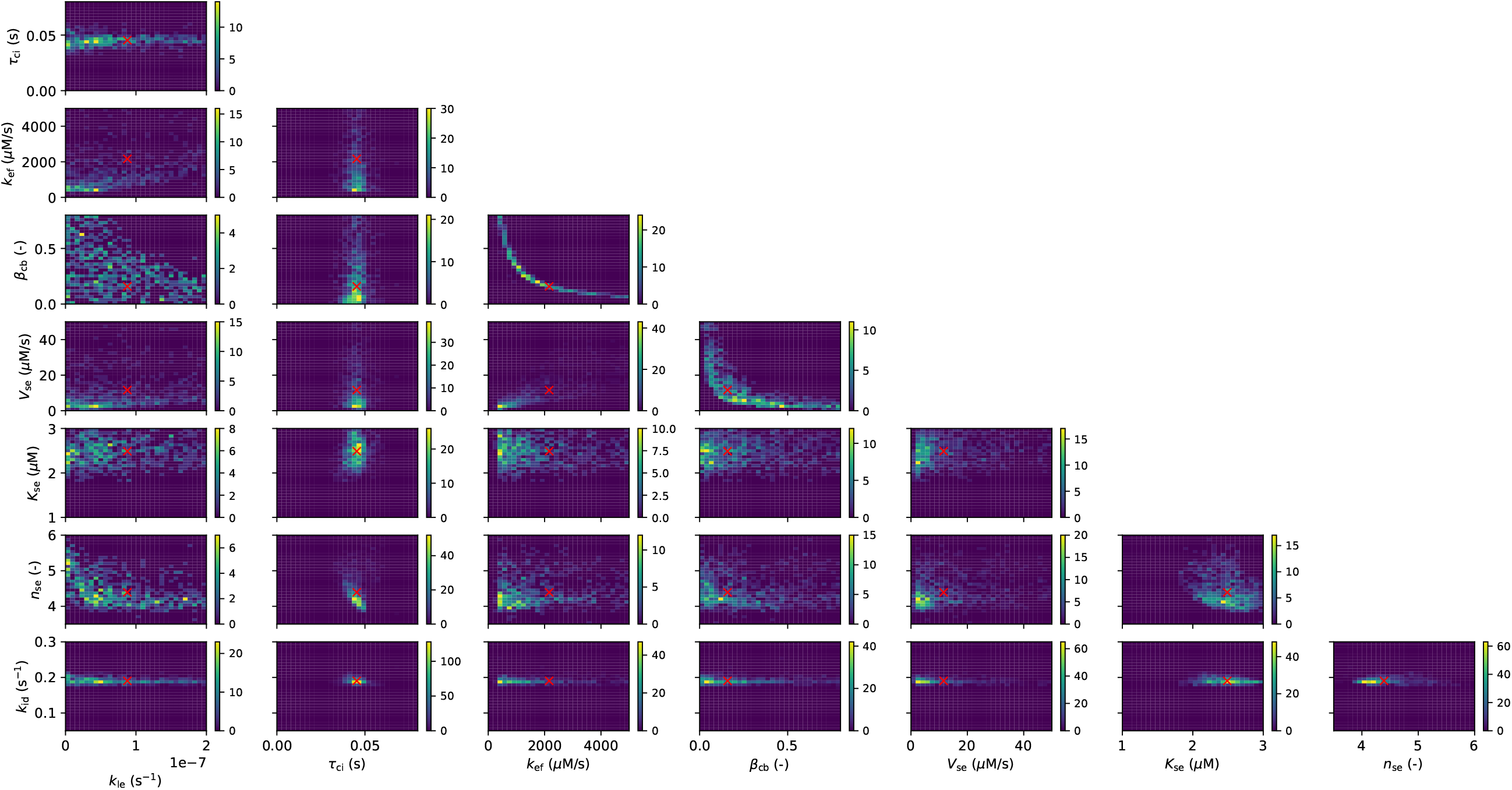
Density parameter space for the Baseline model (presented as 1 parameter vs 1 parameter). In each subplot the red cross indicates the geometric mean.

EC cytosolic Ca^2+^ transient is known to depend on the Ca^2+^ inactivation time constant (*τ*_ci_) and the IP_3_ dissociation rate (*k*_id_), which for the Baseline model appear to fall in well defined range of values.

The predictive capacity of the Baseline model is also assessed by considering another Ca^2+^ transient feature, the First decline rate, which reflects the rate of Ca^2+^ removal from the cytosol after stimulation. Carter and Ogden [29] extracted this feature from experimental time recordings of Ca^2+^ concentration for different IP_3_ stimulation levels and reported it as function of Ca^2+^ Peak. This feature appears extremely heterogeneous across the EC population and therefore the extracted average trend can be considered less representative than the other population features. Figure 5 shows a comparison between the model predictions and experimentally reported values. Despite its simplicity, the Baseline model can reproduce, from a quantitative point of view, values in line with the main trend of the experimentally recorded traces. In the same figure we report the Pearson correlation coefficient between this feature and the model parameters, evaluated for each stimulation level. As expected, the parameter governing the Ca^2+^ sequestration process *K*_se_ stands out, indicating that pumps with high sensitivity with respect to cytosolic Ca^2+^ level (low *K*_se_) are responsible for sharp Ca^2+^ concentration drops. Altogether these results indicate that the Baseline model is able to capture the mean population features of the considered Ca^2+^ dynamics, constituting a first reference model.

**Figure 5:**
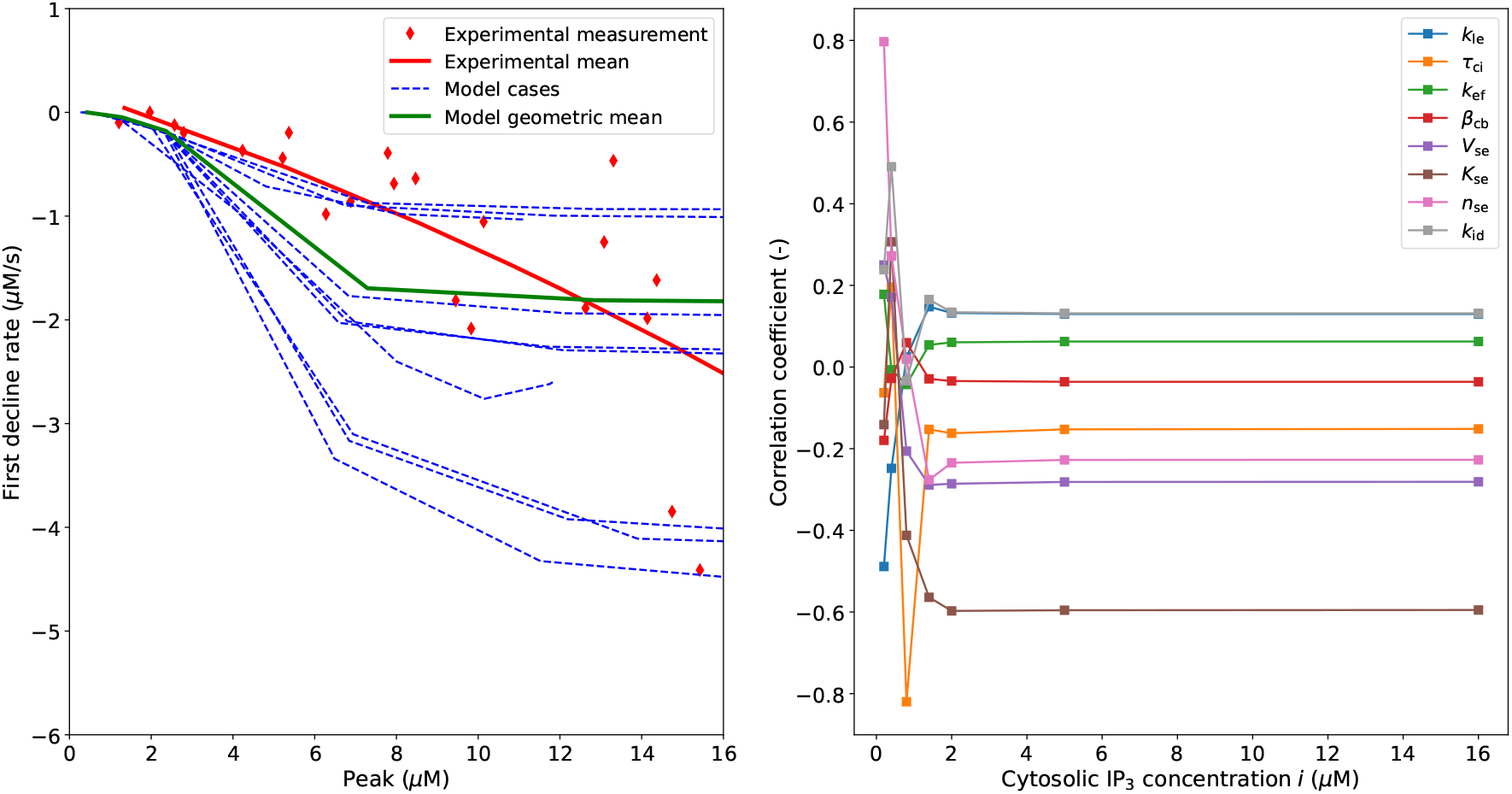
Baseline model validation for 10 model cases/parameter sets and the geometric mean and Pearson correlation coefficient between simulated feature (First decline rate) and parameters.

### 3.2 Ca^2+^ effect on IP_3_Rs activation

The Ca^2+^ dynamics under analysis is initiated by an IP_3_-induced stimulation of the intracellular Ca^2+^ stores. Here we numerically investigate how the dependency of IP_3_-induced ER Ca^2+^ release on cytosolic Ca^2+^ affects the overall EC dynamics. For doing this we consider, with respect to the baseline model, the following variants i) ‘variable’ Ca^2+^ inactivation component (Civar) and ii) ‘variable’ Ca^2+^ inactivation and activation components (Ci&cavar). The parameters of these Ca^2+^-dependent functions are included into the model parameter set to be identified through the optimization algorithm (as in Section 3.1). For only a fraction of the cases (672/1000 for Civar, 642/1000 for Ci&cavar) a solution for the optimization problem was found (cost function below the threshold). The fittings for these two model variants against experimental curves and their parameter spaces are reported in the Appendix (see Figures 12 and 15, and Figures 13 and 16, respectively). With respect to the Baseline model, the variant Civar provides a slightly better estimate in terms of averaged First decline rate (see Figure 14). Table 4 reports the geometric mean (geometric standard deviation) of the parameter set cases across the considered model variants. For most of the simulated cases of this model variant, the sensitivity (*K*_ci_) of the inactivation function seems to be just below 1.0 *μ*M (see Figures 13 and 16 in the Appendix), whilst the distribution of the exponent of the Hill equation (*n*_ci_) appears to be more sparse. On the contrary, a function for ER Ca^2+^ release which accounts for both Ca^2+^-activation and -inactivation effects (Ci&cavar) does not improve the discrepancy with respect to experimental recordings (fitting and validation in the Appendix, respectively Figures 15 and 17). The Ca^2+^ inactivation time constant (*τ*_ci_) does not significantly change across the considered models while the IP_3_ dissociation rate (*k*_id_) appears to be reduced for the two more complex variants.

**Table 4:** Geometric mean (geometric standard deviation) of the parameter set cases for the considered model variants. ^*^: value imposed and not obtained from fitting.

|  | Baseline<br>(847 cases) | Civar<br>(672 cases) | Ci&cavar<br>(642 cases) |
| --- | --- | --- | --- |
| $k_{le}$ ($s^{-1}$) | 8.8e-8 (5.462) | 7.3e-8 (5.741) | 7.1e-8 (5.355) |
| $\tau_{ci}$ (s) | 4.535e-2 (1.247) | 4.874e-2 (1.282) | 5.018e-2 (1.555) |
| $k_{ef}$ ($\mu M/s$) | 2.162e3 (3.580) | 2.167e3 (3.491) | 2.265e3 (3.036) |
| $\beta_{cb}$ (-) | 1.586e-1 (3.553) | 1.559e-1 (3.445) | 1.668e-1 (2.953) |
| $V_{se}$ ($\mu M/s$) | 1.159e1 (3.954) | 1.172e1 (3.973) | 1.116e1 (3.920) |
| $K_{se}$ ($\mu M$) | 2.489 (1.166) | 1.894 (1.253) | 1.916 (1.436) |
| $n_{se}$ (-) | 4.392 (1.101) | 4.802 (1.175) | 4.837 (1.180) |
| $k_{id}$ ($s^{-1}$) | 1.904e-1 (1.068) | 1.516e-1 (1.165) | 1.498e-1 (1.155) |
| $K_{ci}$ ($\mu M$) | 1.0* | 7.630e-1 (1.235) | 7.759e-1 (1.170) |
| $n_{ci}$ (-) | 3.8* | 4.851 (2.146) | 5.125 (2.260) |
| $K_{ca}$ ($\mu M$) | - | - | 1.409e-1 (8.363) |
| $n_{ca}$ (-) | - | - | 2.597 (1.655e1) |

The Ca^2+^-dependency of the IP_3_-induced ER Ca^2+^ release was experimentally assessed by Carter and Ogden by considering different pre-stimulation cytosolic Ca^2+^ levels. The predictions from the Baseline model and the other variants (obtained from the previous fittings) are compared against these experimental data to identify the most appropriate relationship between IP_3_-induced ER Ca^2+^ release and cytosolic Ca^2+^ concentration (see Figure 6). The Baseline and Civar models, despite overestimating the ‘Normalized max flux’ for cytosolic Ca^2+^ concentration below 1.0 *μ*M, are able to capture the main trend of the experimental data. On the other hand, the introduction of a biphasic function accounting for the Ca^2+^ activation component (Ci&cavar) leads to very similar results. We conclude that the inclusion of the Ca^2+^-dependent activation component into ER Ca^2+^ release model does not improve its predictive capacity (and therefore it is discarded in the following) while the Baseline model can be considered representative of the model Civar.

**Figure 6:**
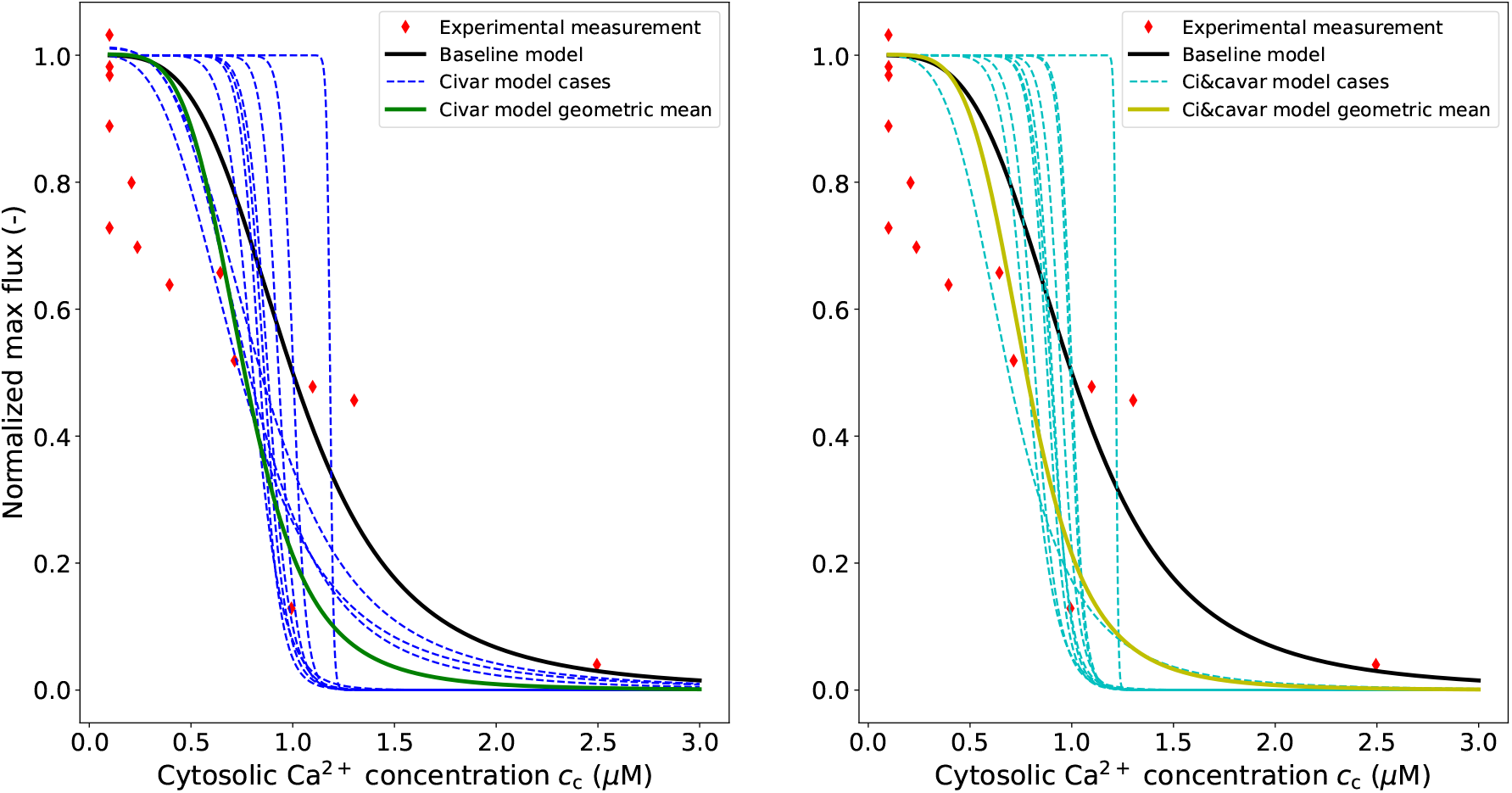
Role of Ca^2+^ on IP_3_Rs function: experimental vs numerical data. In the experiment by Carter and Ogden [29], the cytosolic Ca^2+^ concentration was increased by membrane hyperpolarization before agonist intervention. The ‘Normalized max flux’ was evaluated as a ratio between the Max flux obtained at the imposed (through membrane hyperpolarization) cytosolic Ca^2+^ concentration and the Max flux obtained at basal (before membrane hyperpolarization) cytosolic Ca^2+^ concentration.

### 3.3 Role of cytosolic Ca^2+^ buffering on Ca^2+^ elevation

Here we investigate the role of cytosolic Ca^2+^ buffering on the model’s dynamics. Two variants, with respect to the Baseline model, are considered: i) the buffering parameter *β*_cb_ depends via Eq. (9) on cytosolic Ca^2+^ concentration (*β*_cb_var) and ii) buffering considered as a dynamic process (Dynbuff), with the buffered cytosolic Ca^2+^ concentration obtained by solving Eq. (7). Note that in the latter case the cell dynamics is obtained by solving Eq. (7) in conjunction with Eq. (3) and (10). For the variants *β*_cb_var and Dynbuff a solution for the optimization problem was found, respectively, in 774 and 708 (out of 1000) cases (in the Appendix: see Figures 18 and 21 for their fitting against experimental data, see Figures 19 and 22 for their parameter spaces). Table 5 summarises the geometric mean (across the parameter sets) for each model variant of cytosolic Ca^2+^ buffering. Among the considered models, the changes in Ca^2+^ inactivation time constant (*τ*_ci_) and IP_3_ dissociation rate (*k*_id_) are limited. On the other hand, the non-constant buffering effect (associated with the two variants) reduces the necessary Ca^2+^ sequestration and the maximum store Ca^2+^ efflux (*V*_se_ and *k*_ef_ are significantly smaller for the two variants). Figure 7 shows the Ca^2+^ dynamics (through the time-recordings of *c*_c_ and *h*) across the considered models for different buffering conditions (~ 50% nominal parameter value – ~ 200% nominal parameter value). For all approaches, an increase in the bufferin capacity of the cytosolic proteins reduces the Ca^2+^ Peak but it does not significantly alter the secondary recovery phase (after 4 s). Furthermore, the transient of the gating variable *h* is similar across the model variants, indicating that the (modelled) store Ca^2+^-dependent inactivation is not remarkably affected by the adopted buffering approach. In the Baseline model, buffering plays a more limited action in compensating the Ca^2+^ increase after IP_3_Rs activation. In this case, indeed, the Ca^2+^ sequestration flux (*J*_se_) has a much higher peak than in the buffering variants. These results suggest that, upon IP_3_Rs stimulation, two cells with remarkably different Ca^2+^ sequestration capacity can still exhibit a very similar Ca^2+^ response, as long as there is a cytosolic Ca^2+^ buffering compensation.

**Table 5:** Geometric mean of the parameter set cases for the considered model variants.

| - | Baseline<br>(847 cases) | $\beta_{cb} \text{ var}$<br>(774 cases) | Dynbuff<br>(708 cases) |
| --- | --- | --- | --- |
| $k_{le} \text{ (s}^{-1}\text{)}$ | 8.779e-8 (5.462) | 2.704e-8 (3.842) | 3.488e-8 (3.737) |
| $\tau_{ci} \text{ (s)}$ | 4.535e-2 (1.247) | 4.509e-2 (1.184) | 4.406e-2 (1.459) |
| $k_{ef} \text{ (}\mu\text{M/s)}$ | 2.162e3 (3.580) | 5.874e2 (2.302) | 6.310e2 (2.397) |
| $\beta_{cb} \text{ (-)}$ | 1.586e-1 (3.553) | - | - |
| $k_{bd}^* \text{ (}\mu\text{M)}$ | - | 6.517e2 (1.372e1) | - |
| $k_{bd} \text{ (s}^{-1}\text{)}$ | - | - | 1.155e3 (1.689e1) |
| $V_{se} \text{ (}\mu\text{M/s)}$ | 1.159e1 (3.954) | 3.141 (2.471) | 3.457 (2.856) |
| $K_{se} \text{ (}\mu\text{M)}$ | 2.489 (1.166) | 2.454 (1.141) | 2.540 (1.242) |
| $n_{se} \text{ (-)}$ | 4.392 (1.101) | 4.438 (1.100) | 4.315 (1.124) |
| $k_{id} \text{ (s}^{-1}\text{)}$ | 1.904e-1 (1.068) | 1.882e-1 (1.038) | 1.895e-1 (1.048) |
| $B_T \text{ (}\mu\text{M)}$ | - | 7.943e1 (2.486e1) | 5.589e1 (2.274e1) |
| $k_{ba} \text{ (}\mu\text{M}^{-1}\text{s}^{-1}\text{)}$ | - | - | 1.184e1 (1.546e1) |

**Figure 7:**
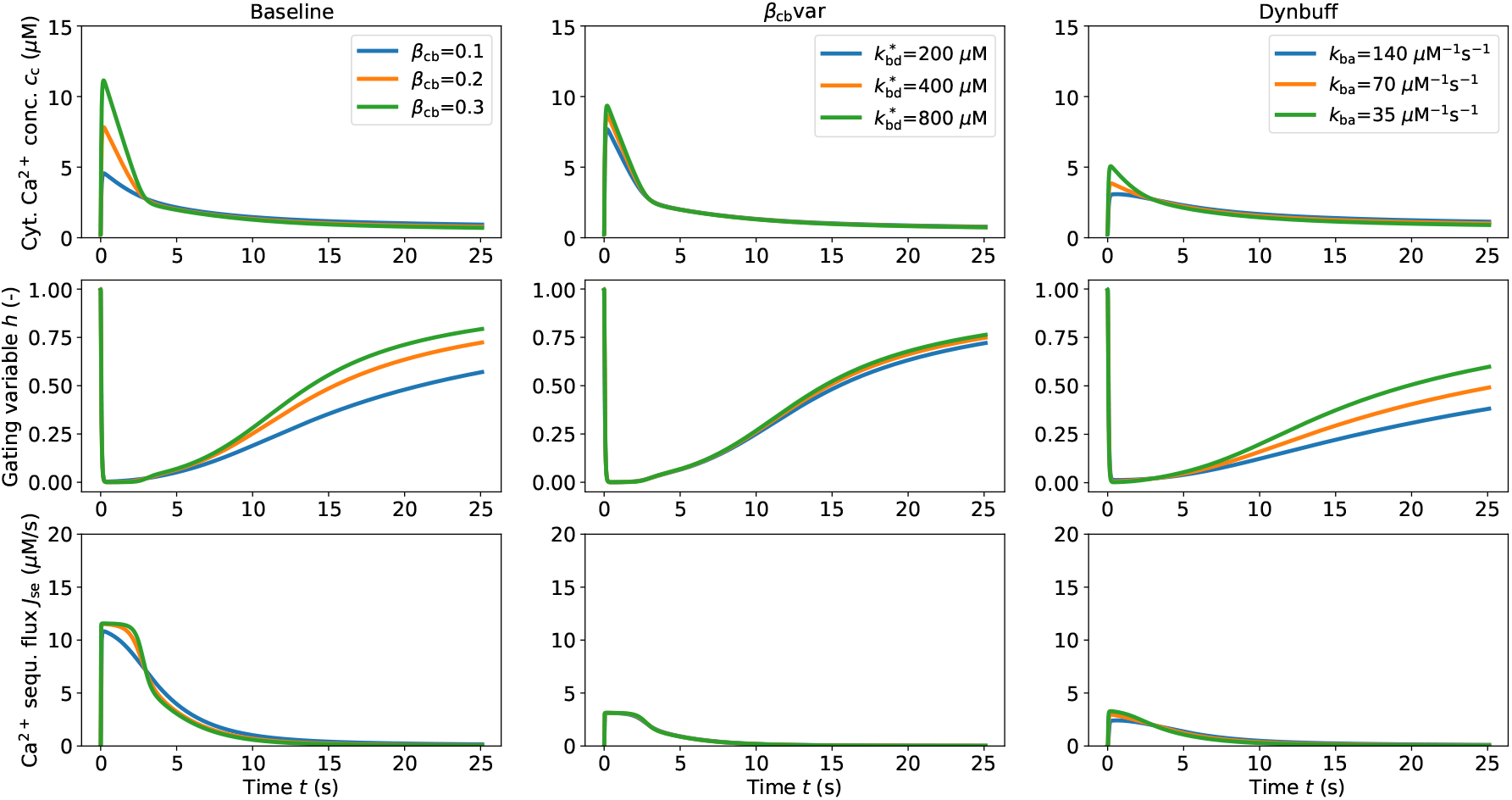
Effects of cytosolic Ca^2+^ buffering on the cell Ca^2+^ dynamics. The reported curves correspond to the geometric mean of the considered model variants with IP_3_ stimulus 1.4*μ*M.

Despite the Ca^2+^ store IP_3_-sensitivity function for porcine aortic EC being well-defined in [29], it is important to assess its role on the overall cell dynamics. We therefore numerically explore, for all the buffering approaches, how the variability in IP_3_Rs’ sensitivity to IP_3_ may affect the EC Ca^2+^ response to different IP_3_ stimulation levels (see Figure 8). As expected, an increase in IP_3_Rs’ sensitivity to IP_3_ (corresponding to a *K*_ia_ reduction) makes the cell more responsive to low concentration of stimulating agent. The Ca^2+^ peaks obtained with IP_3_ stimuli 1.4 *μ*M and 10.0 *μ*M are relatively similar (~30% difference) for *K*_ia_=1.0 *μ*M, whilst they are significantly different (>100% difference) for higher *K*_ia_. Also in this case, no substantial changes in the Ca^2+^ transients are recorded between the different buffering approaches. Given the results of the comparison with experimental data, the reported variants (with more sophisticated buffering approach) may also serve as reference model. However, the increased model complexity (larger number of parameters with respect to the Baseline) does not provide a significantly better description of the Ca^2+^ dynamics under analysis.

**Figure 8:**
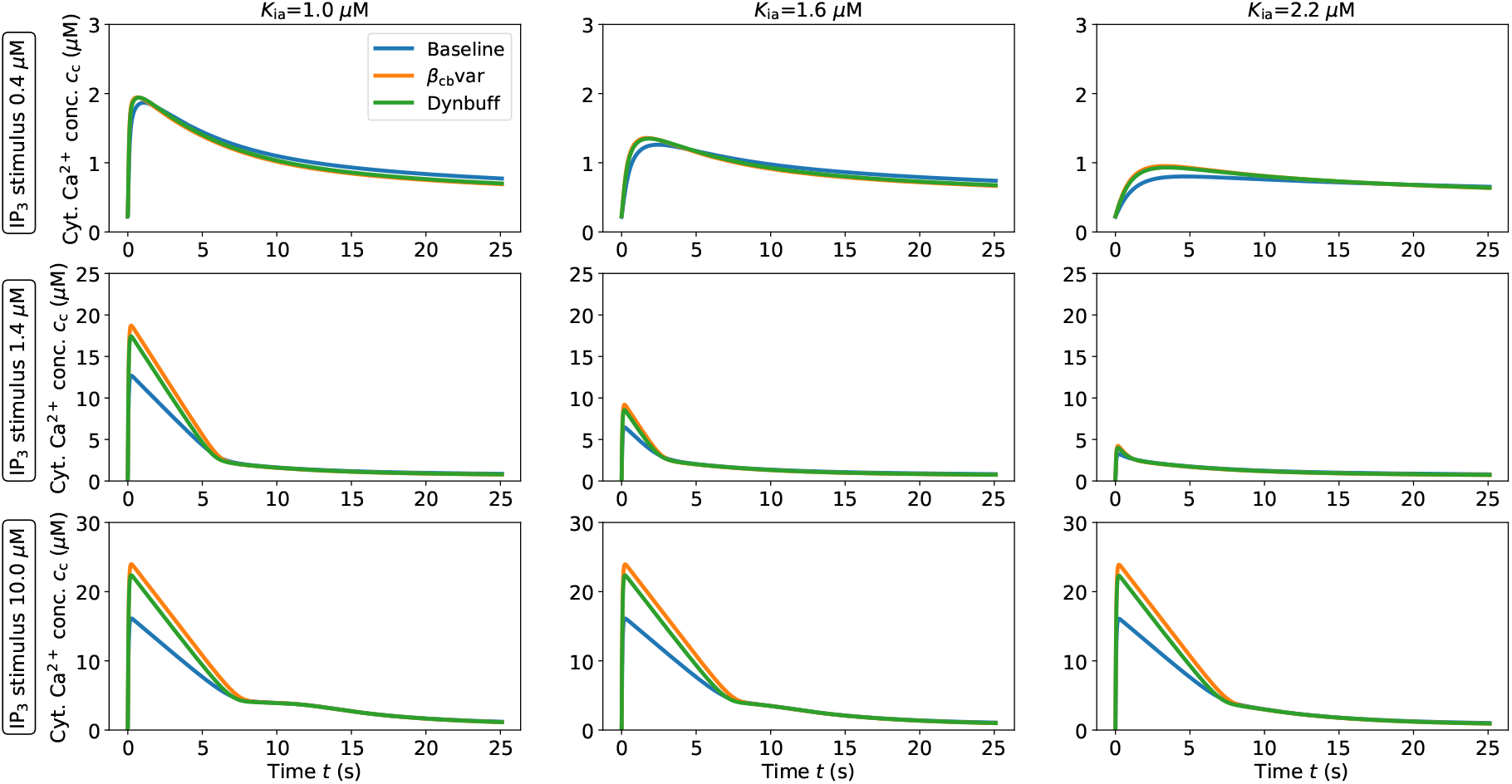
Role of IP_3_Rs’ sensitivity to IP_3_ (*K*_ia_) on the cell cytosolic Ca^2+^ transient. The reported curves correspond to the geometric mean of the considered model variants.

### 3.4 Role of IP_3_ dissociation and Ca^2+^ inactivation time constants on Ca^2+^ dynamics

The IP_3_ dissociation constant (*k*_id_) and the Ca^2+^ inactivation time constant (*τ*_ci_) are expected to play an important role on the cell dynamics. The impact of the former on the Ca^2+^ transient is assessed by means of Figure 9 (for different stimulation levels and buffering approaches). Surprisingly, the IP_3_ rate of dissociation plays an important role on the Ca^2+^ response amplitude (Peak) only upon low IP_3_ stimulation levels. For higher IP_3_ stimuli, the *k*_id_ impacts only the second part of the Ca^2+^ decline/recovery phase whilst the previous part of the transient seems unaffected. Upon high stimulation levels, cytosolic Ca^2+^ concentration decline faster for high *k*_id_ (0.38 s^−1^) which is, from a qualitative point of view, more inline with the experimentally recorded single-cell traces (see Figure 2B-C in [29]) than for the intermediate *k*_id_ case (0.19 s^−1^). Values of *k*_id_ in the range 0.15-0.19 s^−1^ were identified through the parameter optimization procedure as geometric mean for the Baseline model and the other variants as they guarantee a better fit over the whole range of IP_3_ stimulation. However, in case the experimentally-derived mean population Peak [29] was overestimated for low IP_3_ stimulation levels (or a more complex IP_3_ dynamics takes place), a higher IP_3_ dissociation constant would represent a more appropriate choice. While the impact of *k*_id_ on the Ca^2+^ transient seems the same across all buffering approaches, the same cannot be said regarding the parameter *τ*_ci_ (see Figure 10). For the Baseline model and upon medium-high stimulation levels, the Ca^2+^ inactivation time constant can significantly modulate the amplitude of the signal but the other two variants appear almost insensitive with respect to this parameter. Interestingly, for different buffering approaches, almost the same geometric mean of *τ*_ci_ was found (~0.045 s). The Ca^2+^-dependent gating variable (*h*) sharply drops after the stimulation in a similar fashion across all buffering approaches (see Figure 11 in the Appendix). The discrepancy in *τ*_ci_-sensitivity between the Baseline and the other variants may be explained as result of the significant difference in the nominal ionic flux (*k*_ef_). Since the magnitude of *k*_ef_ for the Baseline model is significantly higher than for the variants, a variation in IP_3_Rs inactivation time will have a much bigger impact on the cytosolic Ca^2+^ balance.

**Figure 9:**
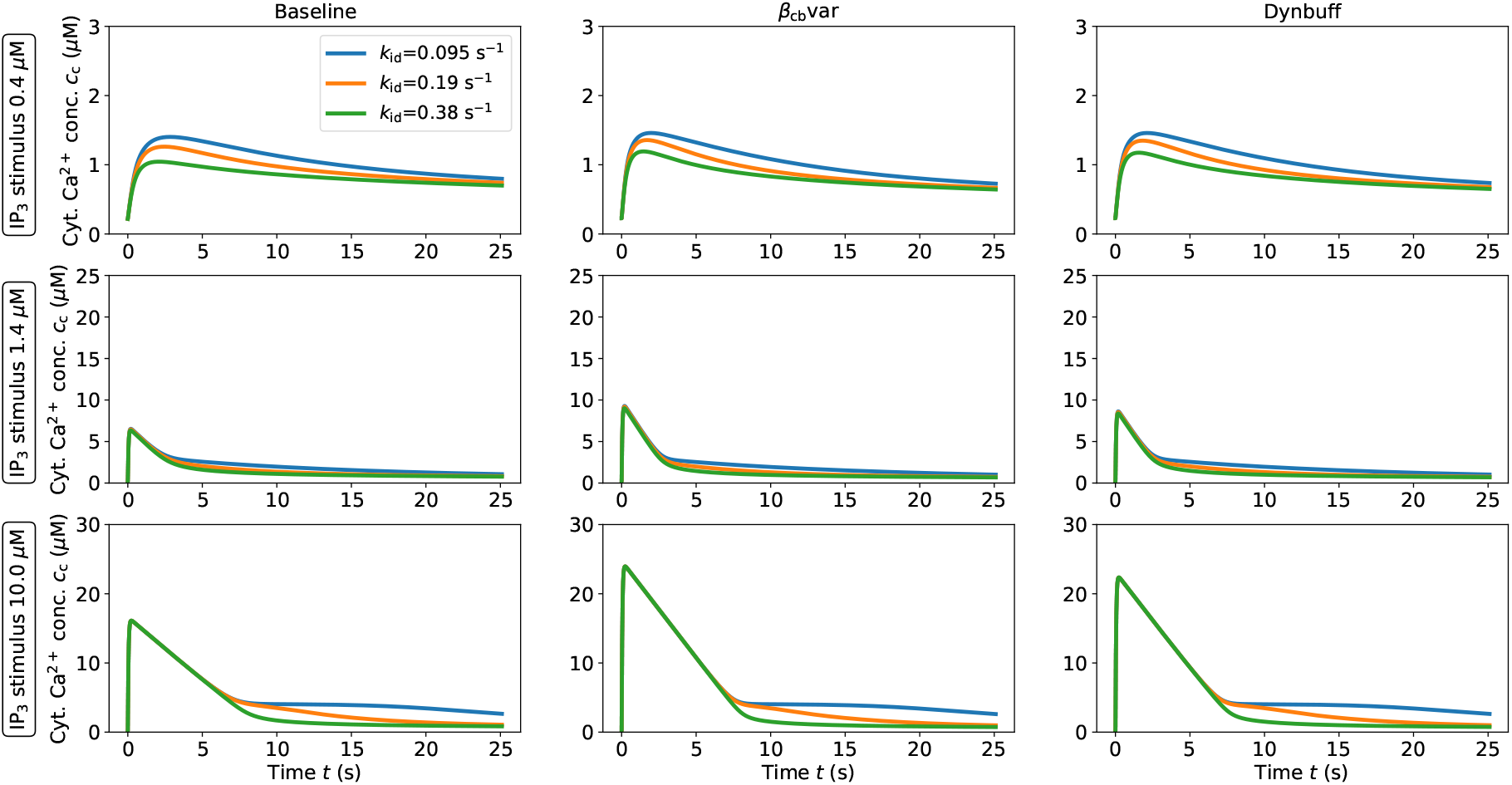
Role of IP_3_ dissociation constant *k*_id_ on the cell cytosolic Ca^2+^ transient. The reported curves correspond to the geometric mean of the considered model variants.

**Figure 10:**
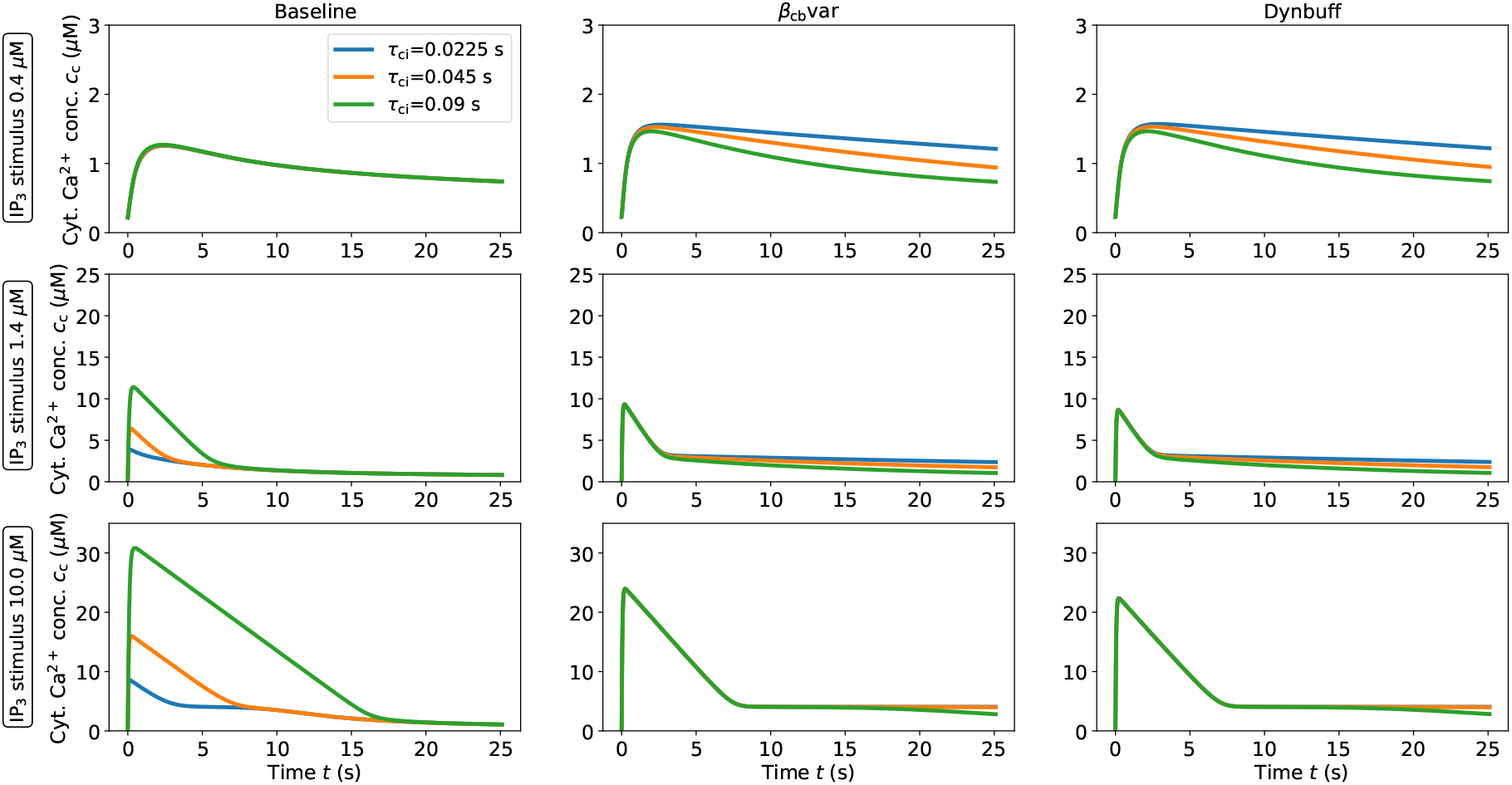
Role of Ca^2+^ inactivation time constant *τ*_ci_ on the cell cytosolic Ca^2+^ transient. The reported curves correspond to the geometric mean of the considered model variants.

**Figure 11:**
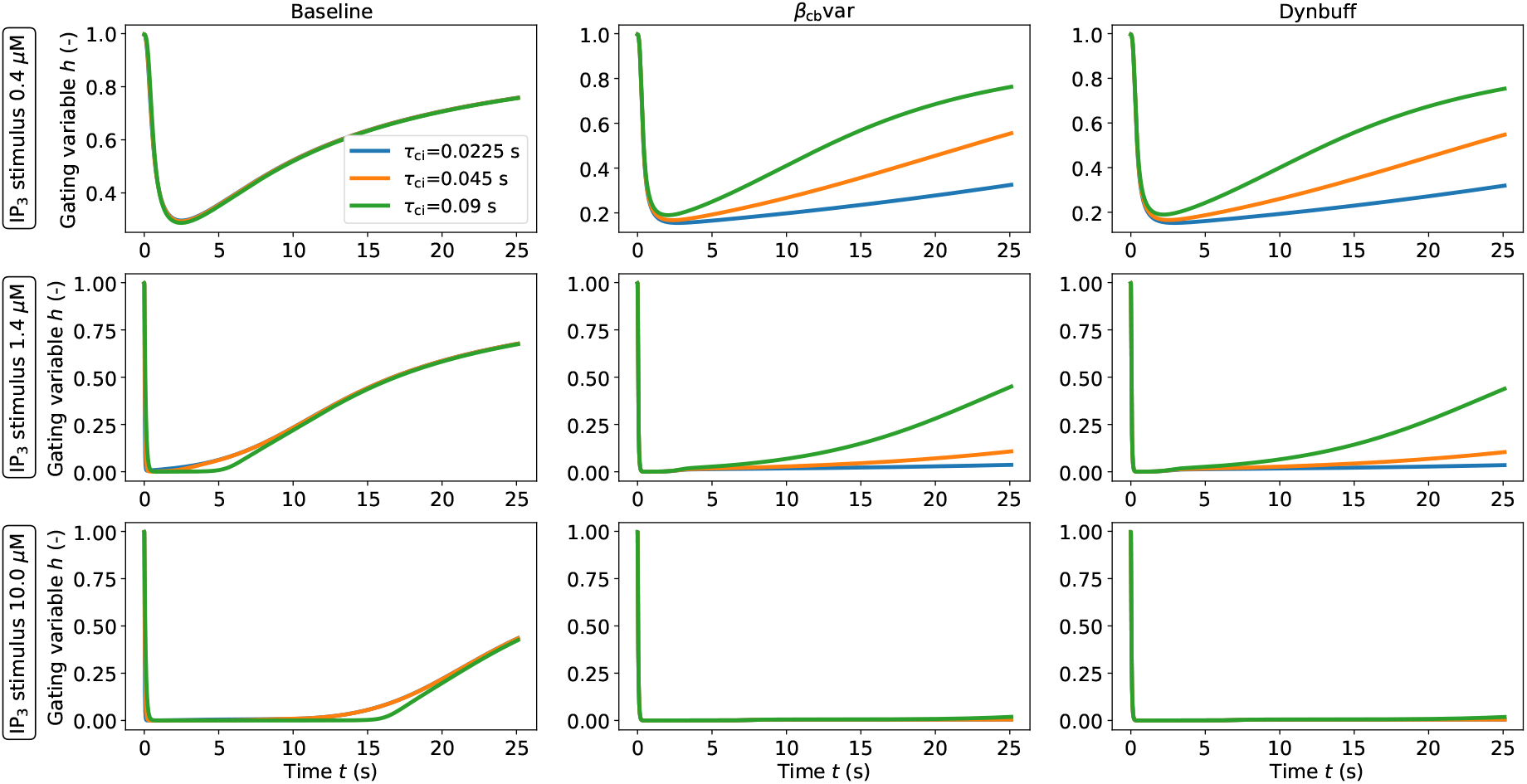
Role of Ca^2+^ inactivation time constant *τ*_ci_ on the Ca^2+^-dependent gating variable transient. The reported curves correspond to the geometric mean of the considered model variants.

**Figure 12:**
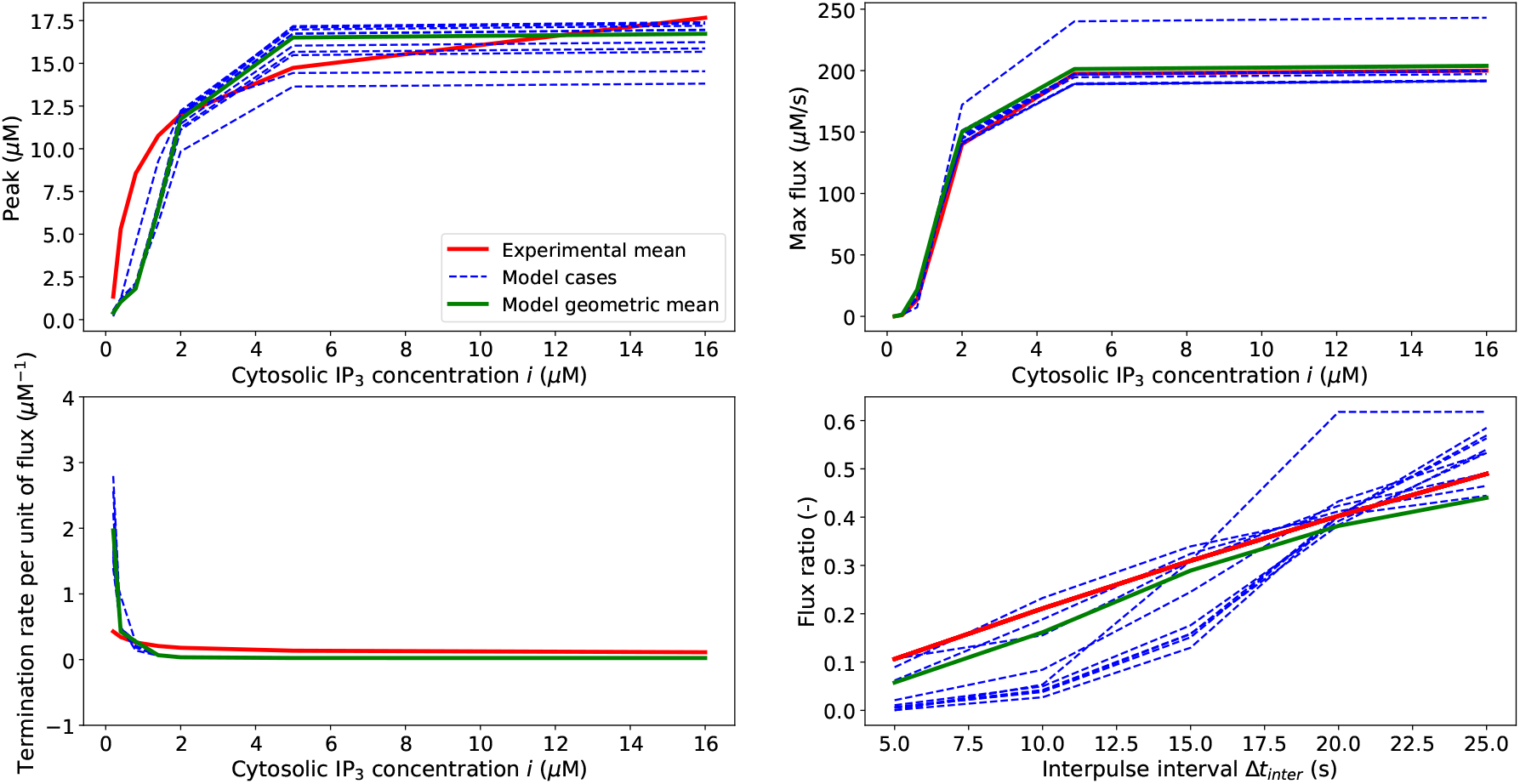
Comparison between the experimental mean against 10 model cases/parameter sets and geometric mean of the Civar model.

**Figure 13:**
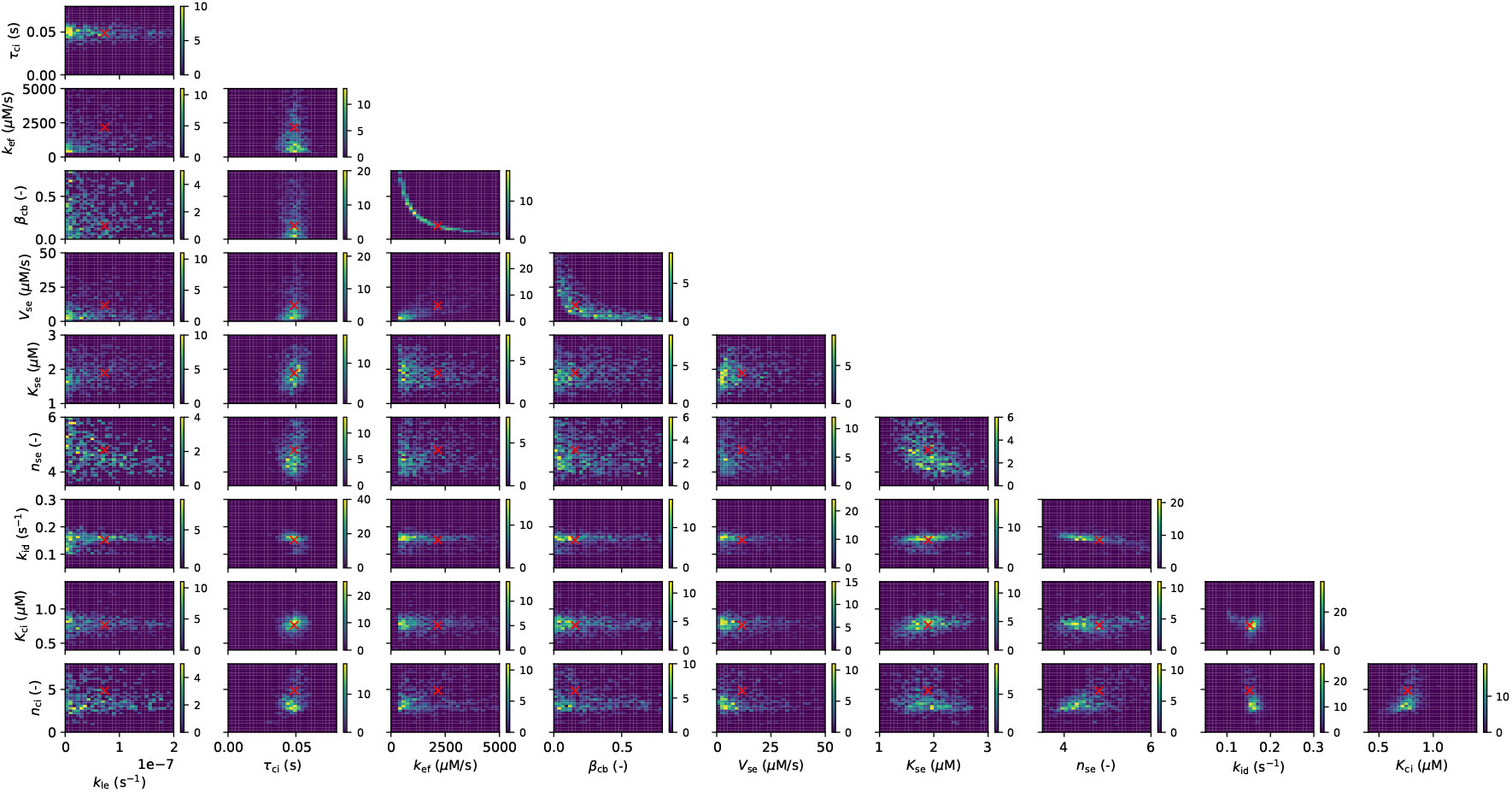
Density parameter space for the Civar model (presented as 1 parameter vs 1 parameter). In each subplot the red cross indicates the geometric mean.

**Figure 14:**
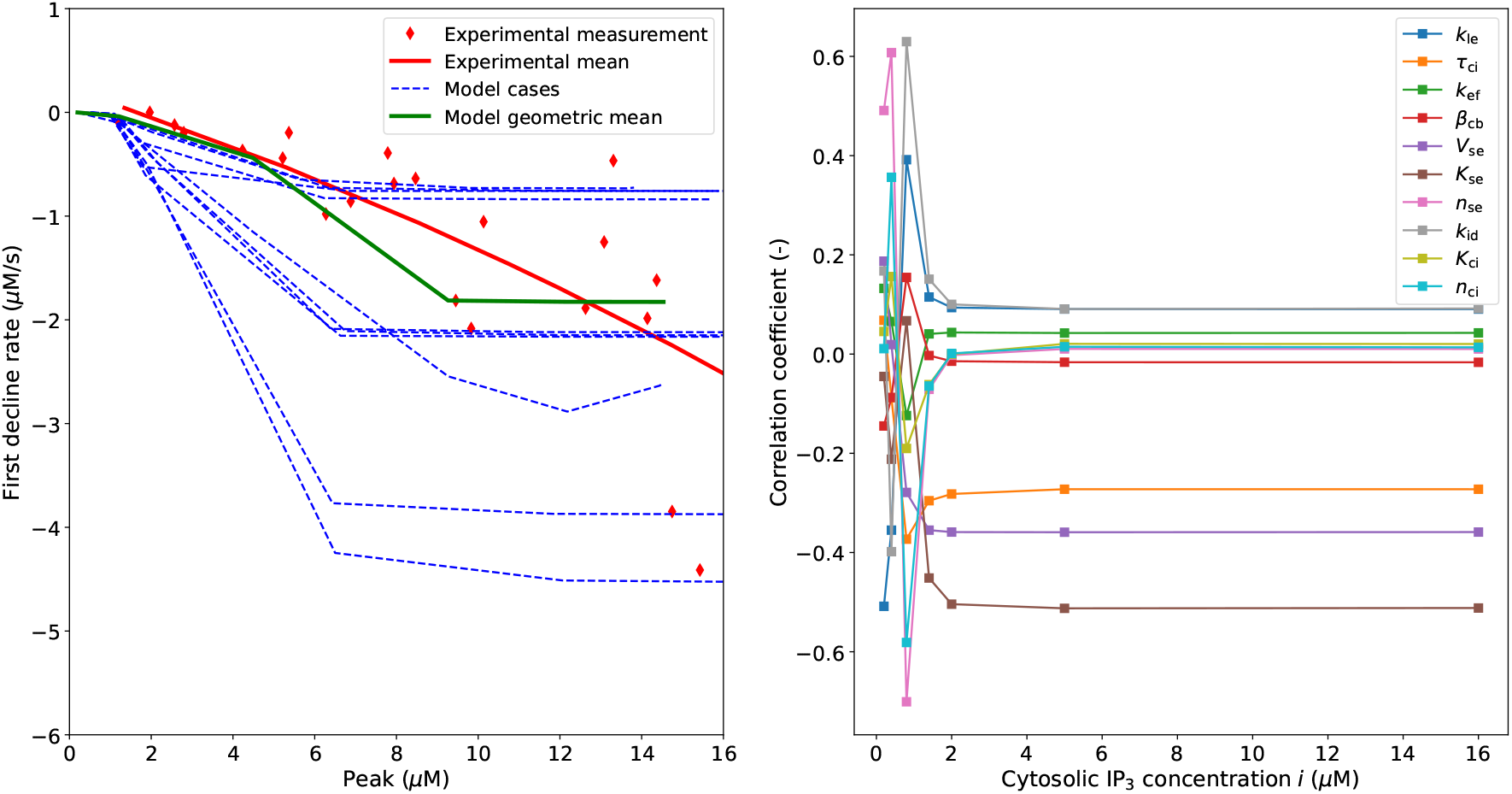
Civar model validation for 10 simulated cases and the geometric mean and Pearson correlation coefficient between simulated feature and parameters.

**Figure 15:**
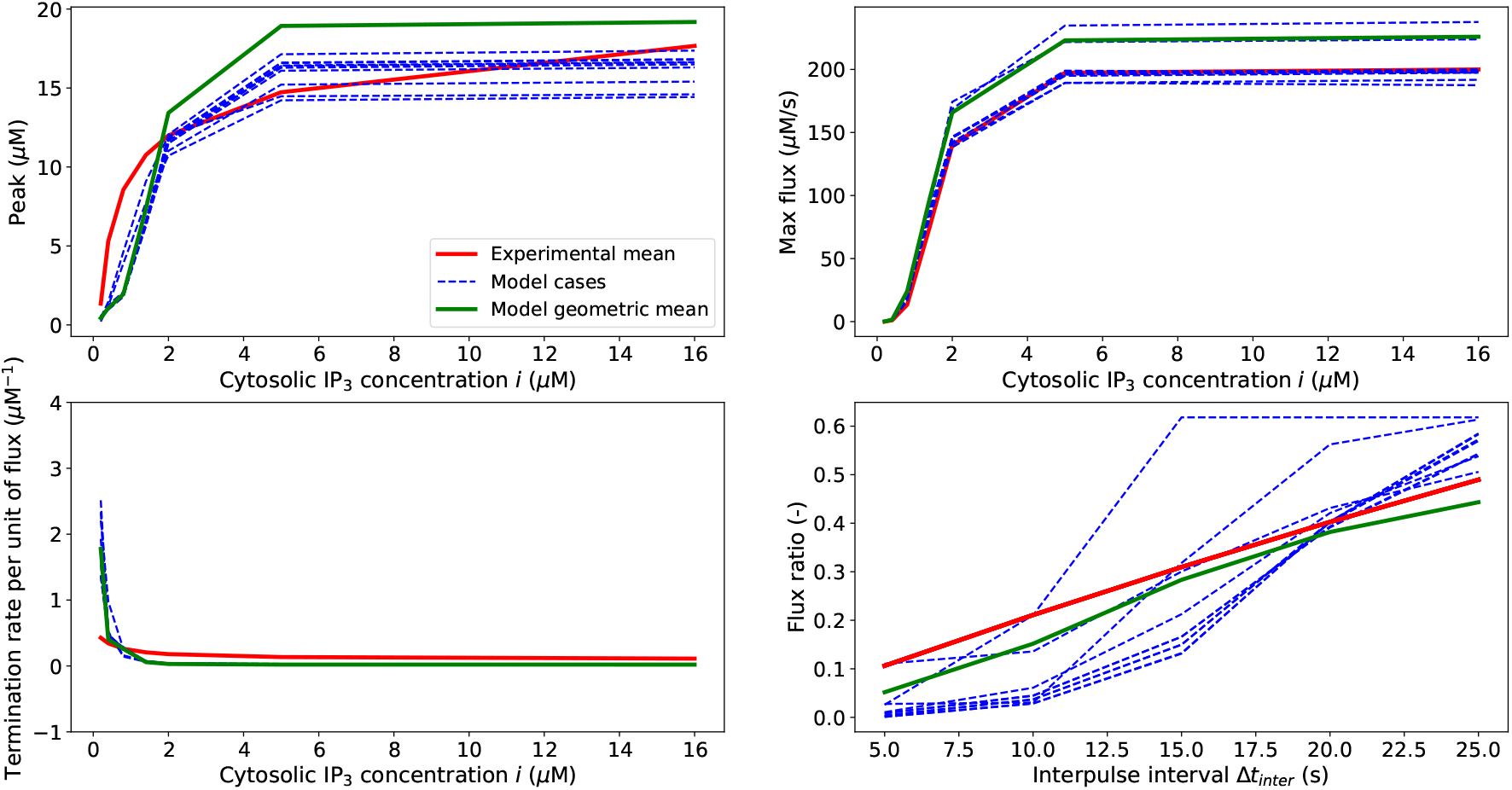
Comparison between the experimental mean against 10 model cases/parameter sets and geometric mean of the Ci&cavar model.

**Figure 16:**
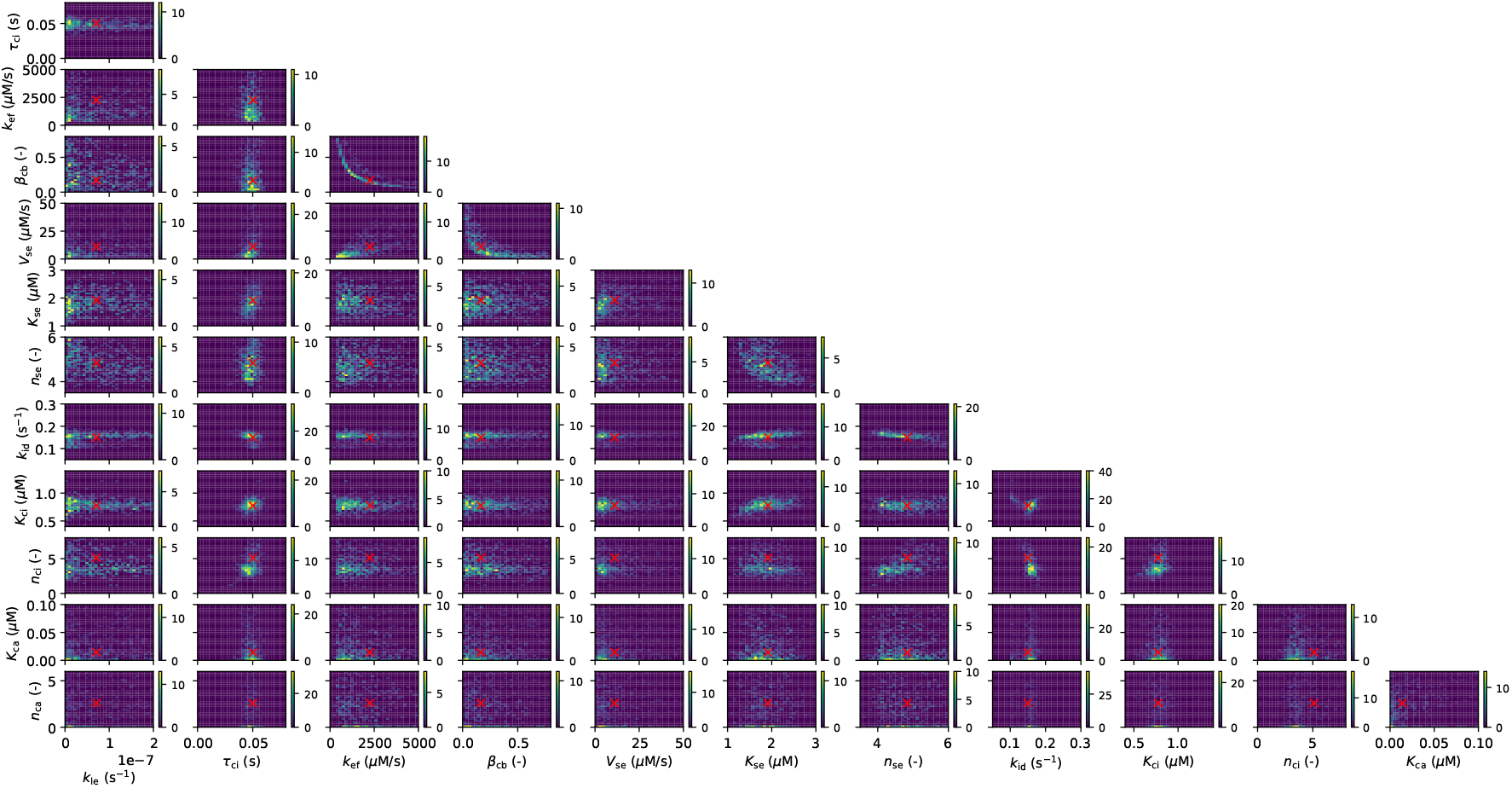
Density parameter space for the Ci&cavar model (presented as 1 parameter vs 1 parameter). In each subplot the red cross indicates the geometric mean.

**Figure 17:**
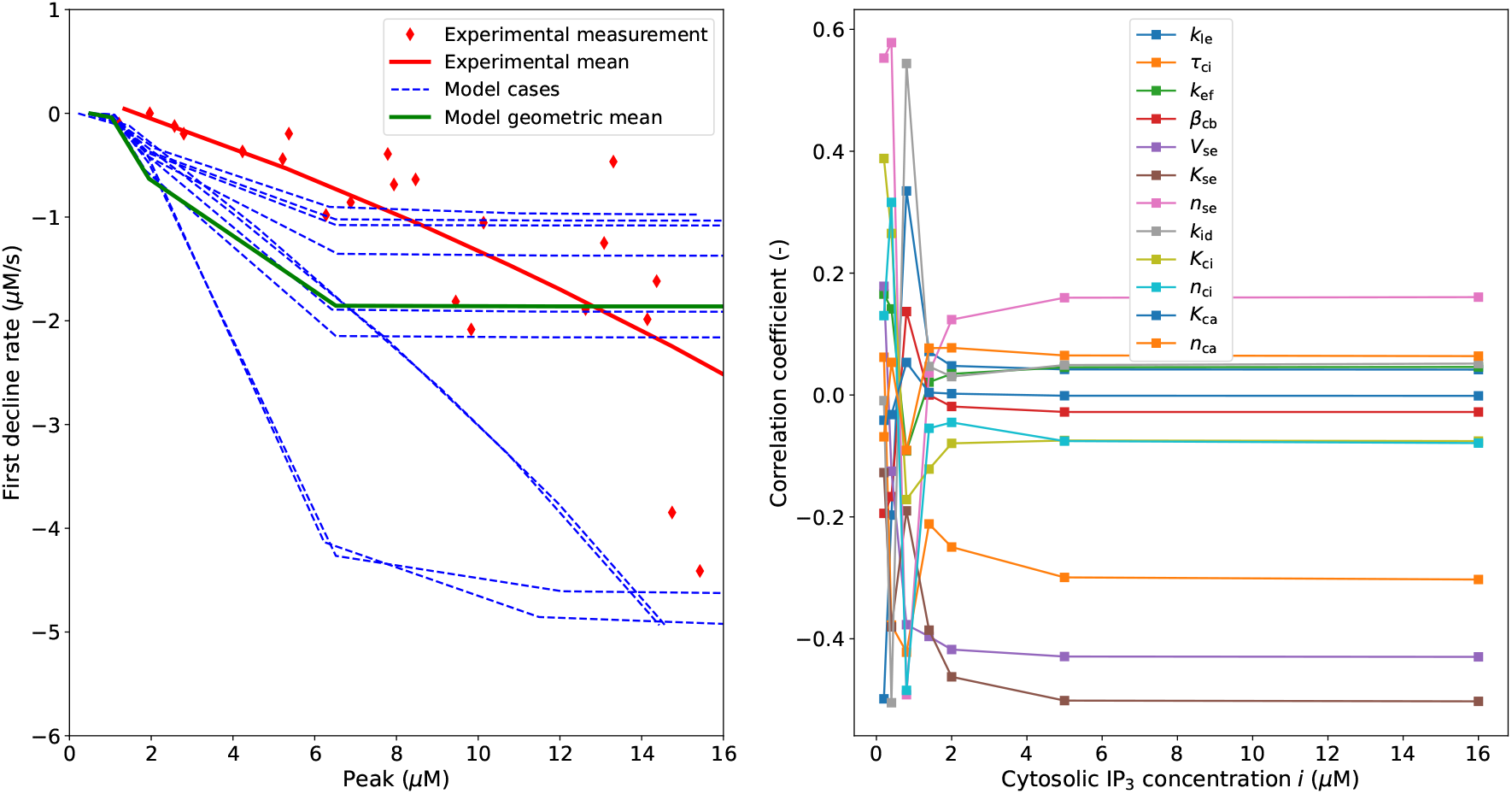
Ci&cavar model validation for 10 simulated cases and the geometric mean and Pearson correlation coefficient between simulated feature and parameters.

**Figure 18:**
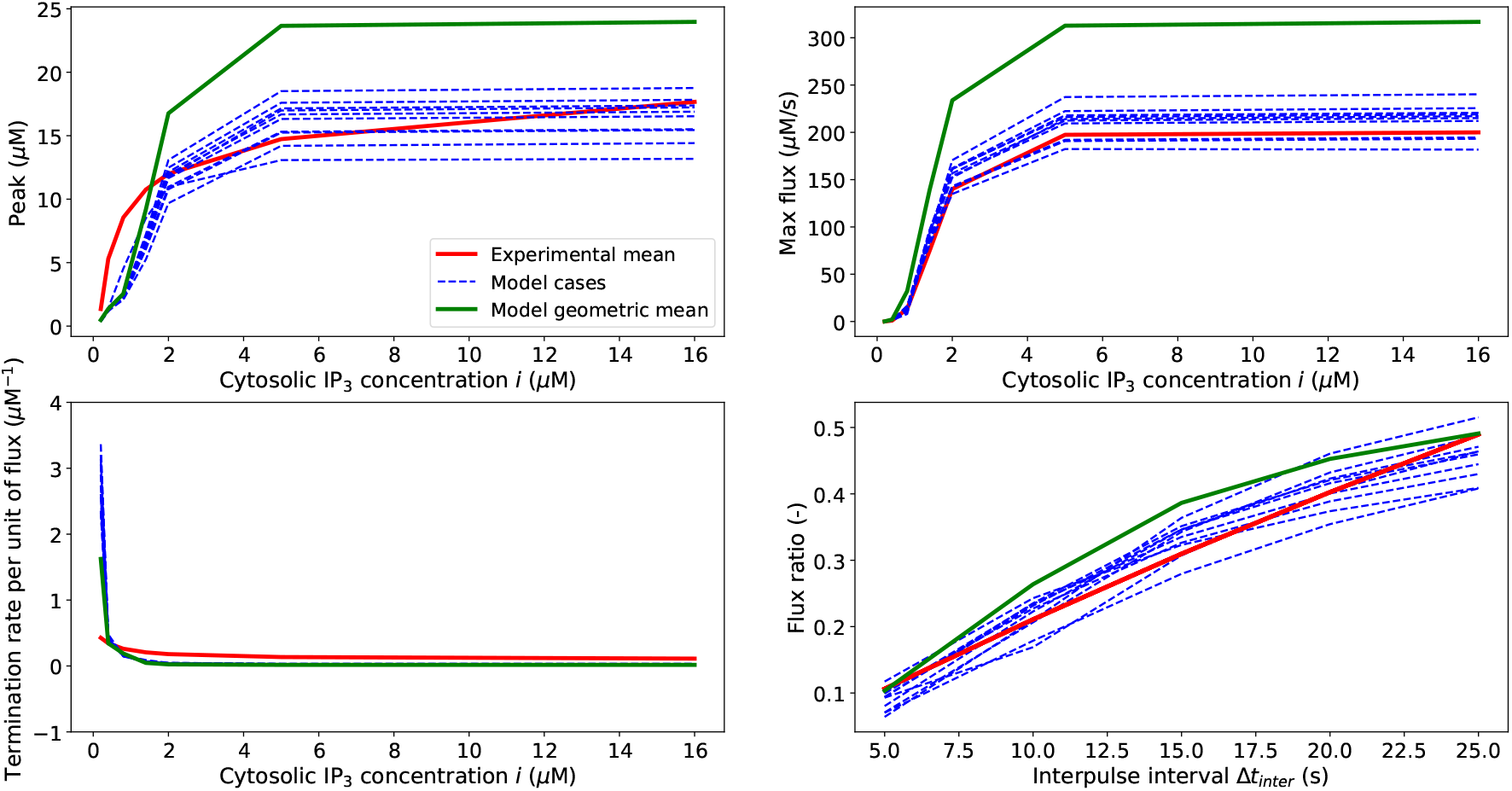
Comparison between the experimental mean against 10 model cases/parameter sets and geometric mean of the *β*_cb_var model.

**Figure 19:**
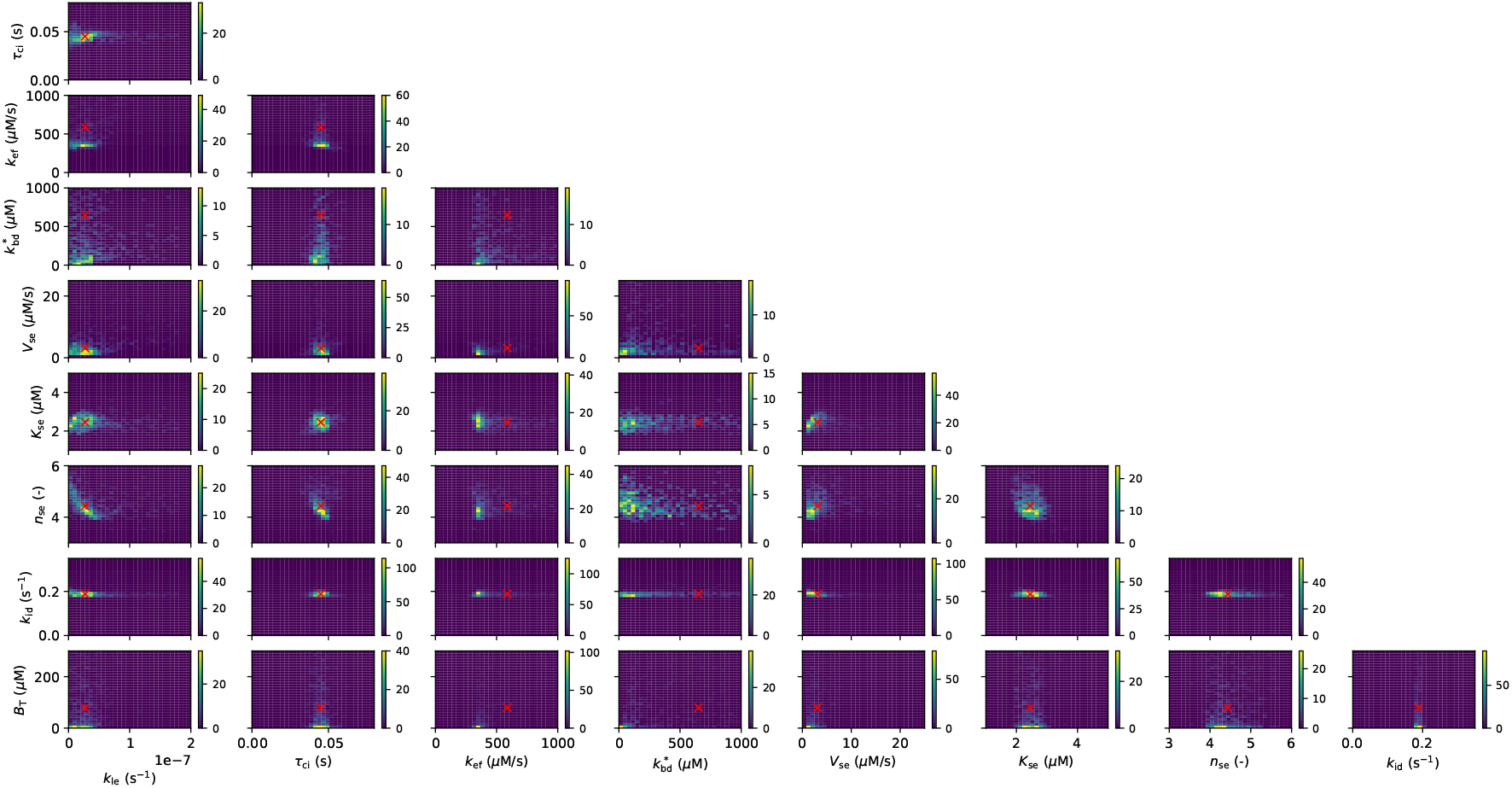
Density parameter space for the *β*_cb_var model (presented as 1 parameter vs 1 parameter). The red cross in the the subplot indicates the geometric mean.

**Figure 20:**
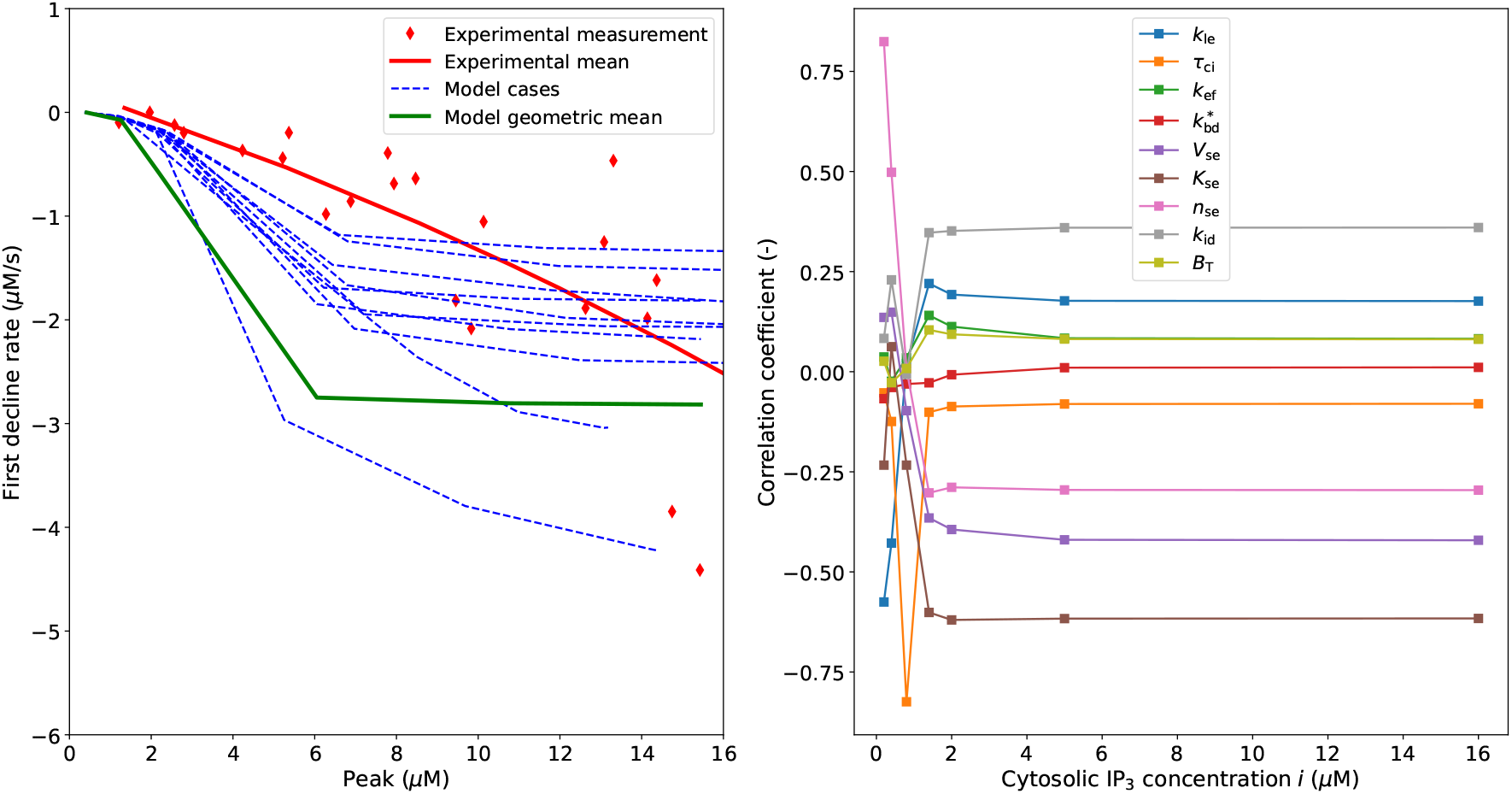
*β*_cb_var model validation for 10 simulated cases and the geometric mean and Pearson correlation coefficient between simulated feature and parameters.

**Figure 21:**
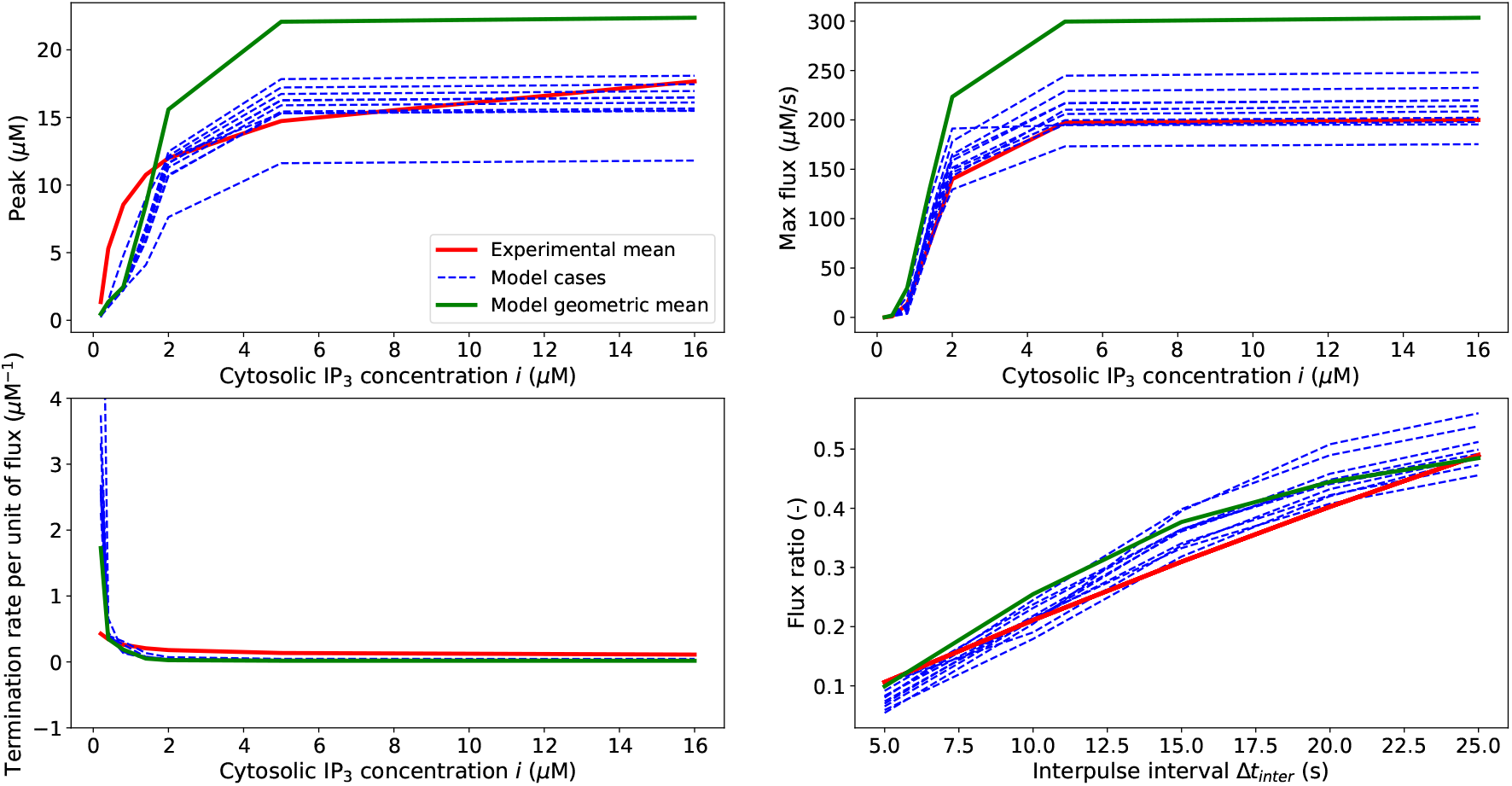
Comparison between the experimental mean against 10 model cases/parameter sets and geometric mean of the Dynbuff model.

**Figure 22:**
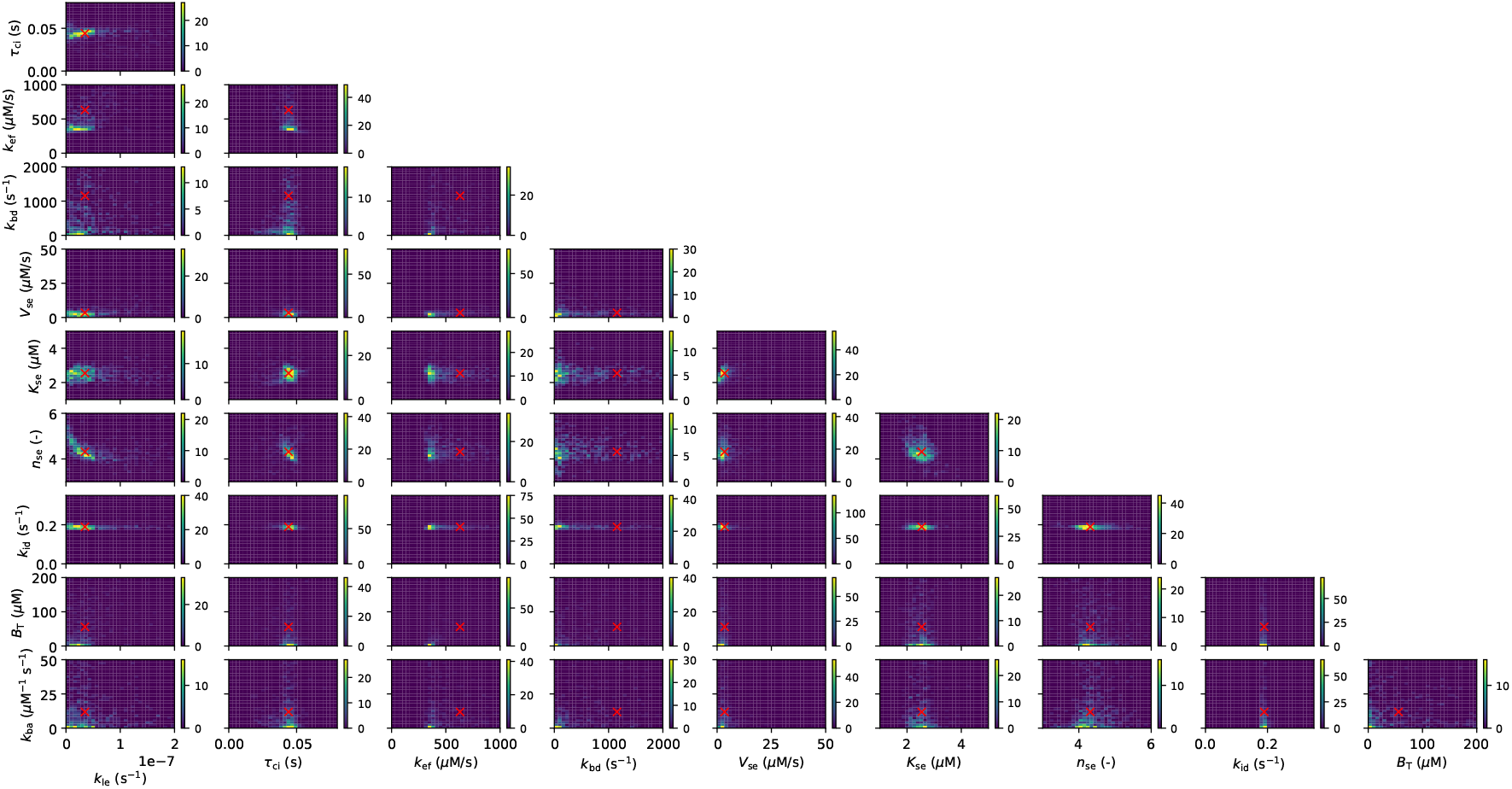
Density parameter space for the Dynbuff model (presented as 1 parameter vs 1 parameter). The red cross in the the subplot indicates the geometric mean.

**Figure 23:**
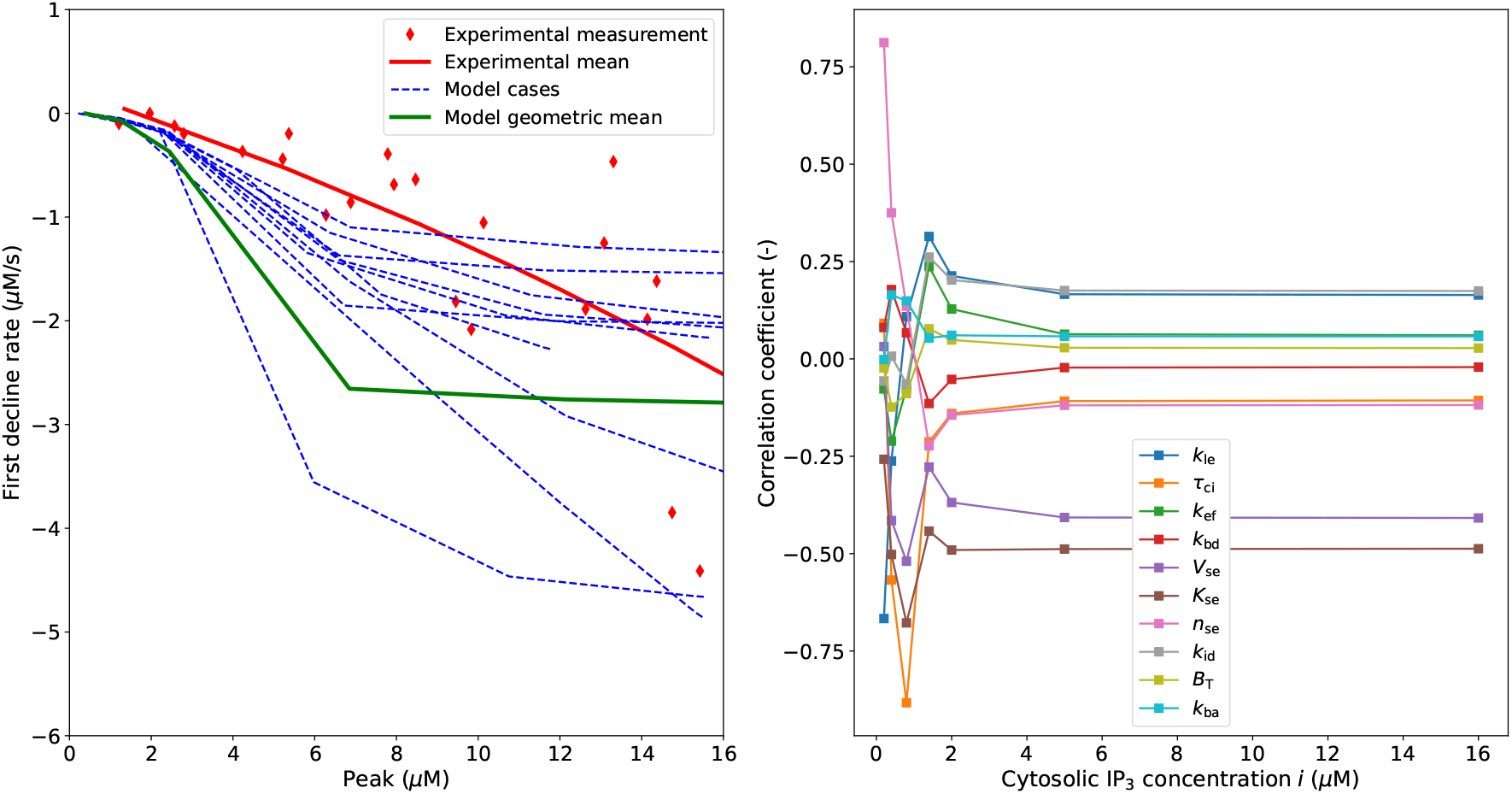
Dynbuff model validation for 10 simulated cases and the geometric mean and Pearson correlation coefficient between simulated feature and parameters.

## 4 Discussion and concluding remarks

Through this work we introduced a minimalistic modelling framework for describing the EC Ca^2+^ dynamics induced by IP_3_Rs activation. For doing this, we hypothesized the existence of a reference model able to the capture the average Ca^2+^ transient features from the cell population used in the study by Carter and Ogden [29]. This experimental protocol enabled the characterization of the Ca^2+^ efflux following the IP_3_Rs activation across a range of stimulation levels. The proposed structure of Baseline model, which considers the cell variables homogeneously distributed in space, allowed a more than satisfactory fitting against four features of the time-recorded Ca^2+^ traces. Furthermore, the validation against the measured First decline rate and Normalized max flux corroborated its predictive capacity. These results together indicate that IP_3_-induced EC dynamics, despite involving diffusive phenomena across the cytosol, can be captured with a good level of accuracy by a lumped modelling approach. Model variants led to similar results under most of the aspects, indicating that the minimalistic Baseline model can constitute a reference framework for capturing the key features of the considered Ca^2+^ dynamics. Whilst variations in the description of IP_3_Rs Ca^2+^-dependent inactivation (Civar) can reduce the discrepancy with respect to measurements, the inclusion of the IP_3_Rs Ca^2+^ activation effect (Ci&cavar) does not substantially improve the IP_3_Rs Ca^2+^-dependent function in terms of accuracy. Supported by findings on other types of cells, more complex mathematical constructs could be used in future for describing the bidirectional link between IP_3_, Ca^2+^ and Ca^2+^ store release. Mak et al. [53] investigated the IP_3_ dependence of the Ca^2+^ sensitivity (of the ER receptors) at single channel level in isolated Xenopus oocyte nuclei by employing a biphasic Hill equation. Whilst the Ca^2+^-activation effect appeared to be independent from *i*, the dissociation constant describing the Ca^2+^ inhibitory role increased with higher *i*. The IP_3_-dependent effect on the Ca^2+^ inhibitory role may be accounted for by defining the parameter *K*_ci_ as dependent (via activation function) on IP_3_ concentration *i*. The inclusion of more sophisticated Ca^2+^ buffering did not significantly alter the shape of Ca^2+^ response, even though the Ca^2+^ sequestration function is reduced. The proposed framework was meant to provide a detailed description of the Ca^2+^ efflux due to IP_3_Rs activation by minimizing the model complexity. For this reason, a very simplified description of the cytosolic Ca^2+^ removal mechanisms was adopted. Their description accounts for a simple dependency on cytosolic Ca^2^+ concentration via activation function and does not take into consideration the delay in the activation of such cell components nor the interaction with other ions. This is expected to affect, at least in part, the shape of the second phase of the Ca^2+^ recovery transient. Considering two separate pumps with different Ca^2+^ sensitivity (*J*_ex_ different from zero) in the model identification did not lead to any significant improvement (results not shown). Since the activation of such cell components affects in a minimal way the Ca^2+^ efflux following the IP_3_Rs activation, its detailed analysis is considered beyond the scope of this work.

Through the proposed approach, we were also able to explore parameters characterizing the IP_3_-induced store Ca^2+^ release. The IP_3_ dissociation constant (*k*_id_) within the EC cytosol obtained from the parameter optimization process appeared to be significantly lower than what estimated/used in other cell models (1.0 s^−1^ in [54], 1.25 s^−1^ in [50], 2.0 s^−1^ in [41]). Here we showed that *k*_id_ affects the peak of the Ca^2+^ transient only for low stimulation levels. As expected, higher values of IP_3_ dissociation rate are associated to a faster cytosolic Ca^2+^ decline, but in this case the discrepancy with the experimental data for low IP_3_ stimuli increases. Despite the evidence for the regulatory role of Ca^2+^ and PLC on the cytosolic IP_3_ dynamics [54], this feedback mechanism is not mentioned in the considered experiment [29] and its role remains uncertain. Furthermore, the manipulation of the system by introducing fluorescent dye and/or altering the electrolytic balance may also have a significant impact on the IP_3_ dissociation rate. To justify the reported results, one could speculate that the lower IP_3_ dissociation constant (with respect to other cell types) may be due to the fact that, since IP_3_ represents in ECs the primary pathway for agonist stimulation, it needs to be metabolized by cytosolic proteins at slower rate to enable its signalling at low concentrations. This might be the case, as well as a more complex IP_3_ dynamics taking place, and therefore future studies are necessary for addressing this question. This study also led to a first estimate of the Ca^2+^-inactivation time constant associated to IP_3_Rs, which appeared to be consistent across all the model variants.

In the considered settings, the membrane potential was kept equal to 0 mV throughout the experiment with the aim to isolate the store Ca^2+^ efflux. Behringer and Segal [55] investigated the controversial role of membrane potential on Ca^2+^ response to ACh in endothelial tubes. They showed that, under submaximal activation of muscarinic receptors, membrane potential modulates Ca^2+^ influx through the plasma membrane channels in accord with the electro-chemical driving force. On the other hand, previous experimental evidence [56] indicated that, for isolated ECs from small resistance vessels, membrane potential does not seem to play a significant role on sustained muscarinic agonist-induced cytosolic Ca^2+^ elevation. In pressurized rat mesenteric arteries, Ca^2+^ influx through TRPV4 channels plays a primary role in EC cytosolic Ca^2+^ elevation [55, 13]. However, despite their activation is associated with mechanisms of Ca^2+^-induced Ca^2+^-release at IP_3_Rs, these do not seem to significantly contribute to ACh-evoked relaxations [13].The complexity and uncertainty associated with the role of membrane potential in the regulation of endothelial cytosolic Ca^2+^ motivated our choice of considering the simplified setting by Carter and Ogden [29] where membrane potential was kept constant equal to 0. Despite IP_3_Rs activity depends primarily on cytosolic IP_3_ and Ca^2+^ levels, it may be affected by interaction with other organelles such as mitochondria [57] and other factors such as oxidative stress [58]. Furthermore, Wilson et al. suggested that also pressure, by altering the cell geometry, can affect the Ca^2+^ diffusive environment near IP_3_ receptor micro-domains, limiting the IP_3_-mediated Ca^2+^ signals as pressure increases. Therefore, future studies investigating the interaction between EC Ca^2+^ store release and other cytosolic and membrane components are warranted.

This work lays the foundation for future experimental and modelling studies targeting the analysis and quantification of Ca^2+^ dynamics in ECs and its effects on blood vessel contractility and permeability. The proposed model could indeed be combined with other models representing SMCs’ activity [59, 60] for quantifying the interaction between functionally different vascular cells, the resulting wall stress level and lumen deformation. These theoretical findings may also contribute to the development of new models for studying the blood-brain barrier function [61]. Furthermore, this work might help in the analysis of Ca^2+^ intercellular communications and Ca^2+^ wave propagation across cellular networks such as in [62]. To conclude, the proposed minimalistic theoretical model aims to be a reference in-silico tool for deciphering vascular function.

## Appendix

## Ca^2+^ transient features extraction

## Experimental data

Here we report the method for extracting the statistics from the datasets reported in [29]. Digitization of data from plot images was carried out by employing the open-source tool ‘WebPlotDigitizer’.

## Peak

Here the dependency between the Ca^2+^ peak and IP_3_ concentration (*i*) was established by fitting the population data reported in Figure 3A in [29] (considered IP_3_ concentration range: 0.2-16.0 *μ*M) with the following function

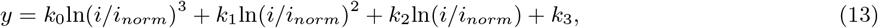

where *i_norm_*=1.0 *μ*M, *k*_0_=6.57e-2 *μ*M, *k*_1_=−0.56 *μ*M, *k*_2_=3.99 *μ*M and *k*_3_=9.48 *μ*M. We highlight that the selected interpolation function leads to a significantly better fitting than linear and quadratic functions.

## Max flux

The population data reporting the maximum Ca^2+^ release from stores against IP_3_ (*i*) concentration was fitted by Carter and Ogden with the following Hill equation (see Figure 3B in [29]):

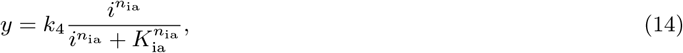

where *k*_4_=200.0 *μ*M/s, *n*_ia_=3.8 and *K*_ia_=1.6 *μ*M. This relationship was obtained by considering IP_3_ stimulation up to 16.0 *μ*M. Beyond this concentration, the max Ca^2+^ release response recorded was extremely variable among cells and therefore was not included.

## Termination rate per unit of flux

Carter and Ogden fitted the measurements of Termination rate per unit flux against IP_3_ concentration (*i*) with a single site binding curve (see Figure 5C in [29])

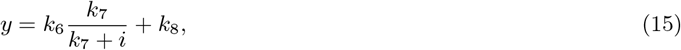

where *k*_6_=0.5 *μ*M^−1^ *k*_7_=0.38 *μ*M and *k*_8_=0.1 *μ* M^−1^.

## Flux ratio

For this feature, the measurements reported in Figure 5C in [29] well-fit with the following function of the interpulse interval (Δ*t_inter_*):

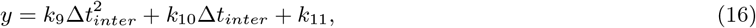

where *k*_9_=−1.1e-4, *k*_10_=2.2e-2 and *k*_11_=−4.3e-3.

## First decline rate

Here the dependency between the first rate of decline and the Ca^2+^ Peak (*Pc*) was established by fitting with a quadratic function the data reported in Figure 3A in [29]

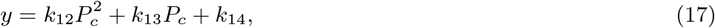

where *k*_12_=−0.003 *μ*M^−1^s^−1^, *k*_13_=−0.127 s^−1^ and *k*_14_=0.218 *μ*M/s.

## Simulated data

Features from the simulated Ca^2+^ transient were extracted as follows. The time point corresponding to the Peak was first identified (by using the Numpy function ‘argmax’), which allowed to calculate the rise time (as peak time minus stimulation time). This allowed to evaluate the time points corresponding to 10 % and 90 % rise time. The Termination rate was evaluated as inverse of the difference between two time points. The Max flux was evaluated as the time derivative of the trace around (averaged over the following 10 time steps) the point corresponding to 10 % rise time. The Termination rate per unit of flux was obtained by dividing the Termination rate by the Max flux. The First decline rate was evaluated as the time derivative of the trace around (averaged over the following 10 time steps) the point corresponding to the lowest time derivative of the trace.

## Role of time constant on Ca^2+^-dependent gating variable

In Figure 11 we report how the Ca^2+^ inactivation time constant *τ*_ci_ affects the transient of the Ca^2+^-dependent gating variable for different buffering approaches.

## Model variants fitting, parameter space and validation

Below the plots of the experimental fitting, parameter space and experimental validation for all the considered model variants (except for the Baseline) are reported.

